# Jetlag Expectations, not Circadian Parameters, Predict Jetlag Symptom Severity in Travelers

**DOI:** 10.1101/2021.04.23.441149

**Authors:** Maximilian Ullrich, Dorothee Fischer, Sebastian Deutsch, Karin Meissner, Eva C Winnebeck

## Abstract

After a flight across multiple time zones, most people show a transient state of circadian misalignment causing temporary malaise known as jetlag disorder. The severity of the elicited symptoms is postulated to depend mostly on circadian factors such as the number of time zones crossed and the direction of travel. Here, we examined the influence of prior expectation on symptom severity, compared to said “classic” determinants, in order to gauge potential psychosocial effects in jetlag disorder.

To this end, we monitored jetlag symptoms in travel-inexperienced individuals (n=90, 18-37y) via detailed questionnaires twice daily for one week before and after flights crossing >3 time zones. We found pronounced differences in individual symptom load that could be grouped into 4 basic symptom trajectories. Both traditional and newly devised metrics of jetlag symptom intensity and duration (accounting for individual symptom trajectories) recapitulated previous results of jetlag prevalence at about 50-60% as well as general symptom dynamics.

Surprisingly, however, regression models showed very low predictive power for any of the jetlag outcomes. The classic circadian determinants, including number of time zones crossed and direction of travel, exhibited little to no link with jetlag symptom intensity and duration. Only expectation emerged as a parameter with systematic, albeit small, predictive value.

These results suggest expectation as a relevant factor in jetlag experience - hinting at potential placebo effects and new treatment options. Our findings also caution against jetlag recommendations based on circadian principles but insufficient evidence linking circadian re-synchronization dynamics with ensuing symptom intensity and duration.

**Significance Statement:** Jetlag disorder afflicts millions of travelers each year - a nuisance on holiday trips but also a danger in safety and performance-critical operations. For effective prevention and treatment, it is critical to understand what influences jetlag severity, i.e. jetlag symptom intensity and duration. In contrast to what guidelines state, in our study, we did not find that symptom severity could be explained by the number of time zones crossed or travel direction. Rather, travelers’ expectations about how long and strongly they will suffer from jetlag symptoms was the only factor systematically predicting jetlag severity. If this holds true not only for subjective but also objective symptoms, we need to revisit assumptions about how circadian desynchronization relates to experienced jetlag symptoms.

## Introduction

Most people become acutely aware that a significant proportion of their physiology is influenced by an endogenous 24-h-timing system, when this circadian system is out of sync with its environment (circadian misalignment). This is the obvious case in jetlag, where the main environmental signal for circadian synchronization, the light-dark cycle, is suddenly shifted in relation to the behavioral rhythm of day and night by flying across multiple time zones. The ensuing symptoms are manifold and cover a wide range of the metabolic, sleep and cognitive spectrum including trouble falling or staying asleep, tiredness, indigestion and problems concentrating ^1-4^.

It is important to note that jetlag, as a term and concept, is used ambiguously across the circadian literature ^5^ to describe three related but distinct phenomena: (i) a rapid shift in the timing of a circadian Zeitgeber signal, (ii) the resulting external and internal circadian misalignment and re-entrainment, as well as (iii) the ultimately manifested downstream symptomatology. The later phenomenon is also termed jetlag disorder by official classifications of the American Academy of Sleep Medicine ^6^ but we shall refer to it under the term ‘syndrome’ to highlight the fact that it is a set of different symptoms and signs. Recommendations for travelers and their physicians on jetlag syndrome are based on evidence synthesized across all three phenomena - without particular distinction between them. They generally state that the severity of jetlag syndrome depends predominantly on the number of time zones crossed, the direction of travel, individual circadian characteristics, light exposure and travel-related factors ^1,7-10^.

Although seldom investigated, it seems entirely plausible that also psychological factors may play a role in the severity of jetlag syndrome. For a multitude of diseases, it is well known that they can be influenced by the psychosocial context such as people’s expectation. Accordingly, for many of the symptoms manifested in jetlag, including gastrointestinal dysregulation, insomnia and tiredness as well as cognitive performance ^11-15^, there is ample evidence outside the context of jetlag that they can be affected by prior expectation. We thus aimed to examine the link between expectation and jetlag syndrome severity as well as the importance of expectation relative to the aforementioned ‘classic’ determinants. Our hope was that delineating the role of expectation in jetlag might open up additional treatment options and ultimately provide further mechanistic insights into the disorder.

However, to examine any effects on jetlag syndrome, jetlag syndrome needs to be quantifiable via a single number or a set of numbers representing its severity, i.e. its duration and/or intensity. Jetlag as the change in Zeitgeber timing (definition i from above) is easily quantified by the shift in light onset or offset as long as photoperiod remains constant e.g. ^16,17^. Quantification of jetlag as circadian re-entrainment (definition ii from above) usually utilizes the time taken for phase resetting as standard measure, based on suitable phase markers such as melatonin, activity rhythms or clock gene expression e.g. ^18,19,20^. In contrast, for jetlag disorder (definition iii from above), there is no gold standard for quantification ^5,7^. Studies have resorted to a number of coarse approaches to gauge jetlag severity based on experienced symptoms. The most direct approach has been simply asking participants for a retrospective rating of their perceived jetlag severity via a single question (“subjective jetlag”) ^20-23^. Some studies employed such a question or assessed selected symptoms on a daily or intra-daily basis, using the results for between-time-point, between-group comparisons but never deriving an overall jetlag intensity or duration metric from this ^24-29^. The only attempt of generating a single jetlag measure from manifested symptoms that we know of was recently undertaken by Becker et al. ^30^. Using their jetlag symptom questionnaire, the Charité Jetlag Scale, the authors determined the highest daily symptom score post-flight and categorized jetlag intensity in relation to values from a control population. Importantly, none of these jetlag measures have so far taken individual symptom dynamics into account, and only a few considered individual pre-flight symptom load ^24,31^. This calls for the development of a finer method of jetlag quantification based on experienced symptoms.

Here, we performed a prospective cohort study in 90 travelers planning flights across >3 time zones to examine potential links between expected and actually experienced jetlag based on manifested symptoms. To this end, we employed a symptom-based definition of jetlag as in jetlag disorder or syndrome (definition iii from above). Given the shortcomings in jetlag quantification from manifested symptoms, we first explored jetlag symptom dynamics pre- and post-flight to develop more granular metrics for jetlag intensity and jetlag duration. Using these new as well as more traditional metrics, we performed regression analyses to identify individual predictors of jetlag and determine the influence of prior expectation.

## Results

Our final study sample consisted of 90 travel-inexperienced volunteers who undertook flights >3 time zones with destinations all around the globe (Tab. 1, Fig. S1A). Up to 1 week before their flight, participants’ status quo was assessed via questionnaires on sociodemographics, health, sleeping patterns and chronotype (Munich ChronoType Questionnaire ^33,34^) as well as jetlag expectation. Following common practice in expectation assessments, we had participants rate their expectation for both symptom intensity and symptom duration. Subsequently, participants recorded the intensity of numerous jetlag symptoms in the morning and evening over 7 days before and after their flight via a detailed questionnaire, the Charité Jetlag Scale ^30^, a validated German adaptation of the two main jetlag questionnaires, the Liverpool and the Columbia Jetlag questionnaires ^31,32^. This provided 2603 complete questionnaires (missingness-matrix in Fig. S1B) and 1308 participant-days for analysis. Demographic composition of our sample, travel details and expectation ratings are listed in Table 1.

**Table 1.**
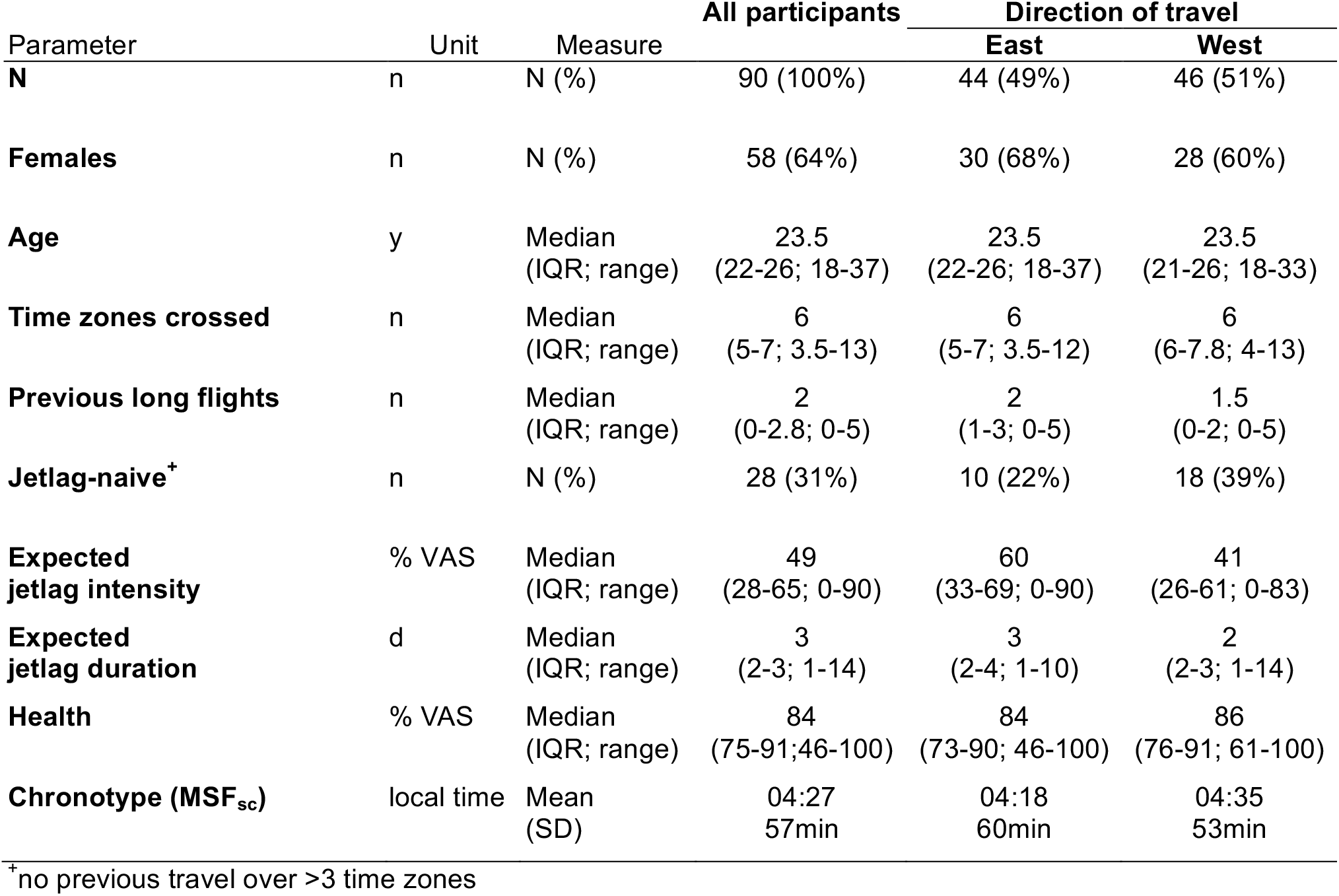
Participant and travel details.

### Substantial inter-individual differences in symptom occurrence and dynamics

The Charité Jetlag Scale provides a total symptom score based on the strength of 15 individual symptoms (from 5 domains across the physiological and cognitive spectrum). A score of 0 indicates no symptoms at all, and each additional point up to the maximum of 60 marks the manifestation of an additional symptom or an increase in the strength of one symptom by 1 unit on a 5-unit scale.

The mean total symptom score over the entire sample showed a trajectory similar to those from previous reports on symptom-based scales ^26,30-32^: Symptom scores were at a low but not asymptomatic baseline before the flight, increased to peak levels in the evening of the first day post-flight – or the flight-day if including this day – and returned to pre-flight levels after about 4 days (Fig. 1A). Quantitative analyses confirmed that symptom load was on average higher after the flight than before (Fig. 1B).

**Figure 1.**
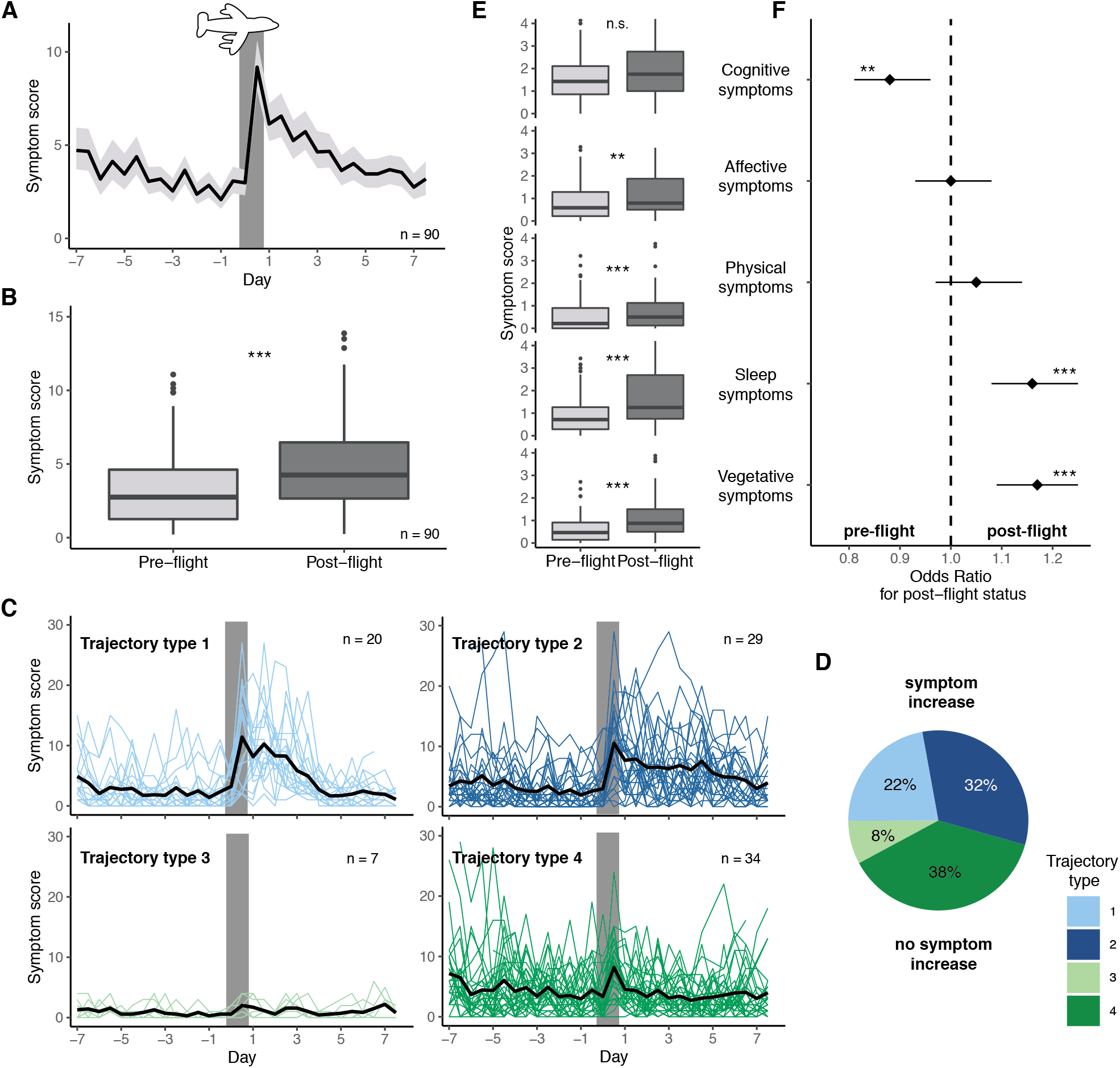
Jetlag symptom dynamics. A) Mean symptom score (± 95% CI) in the morning and evening for 7 days before and after the flight. Grey bar indicates the day of flight. B) Comparison of individual mean symptom scores before the flight (all 7 days) and thereafter (first 4 days). C) Individual symptom scores over the course of the study segregated by trajectory type. Types 1 and 2 show increases in symptom load post-flight, whereas Types 3 and 4 have similar symptom loads throughout the assessment period. Each colored line represents one individual trajectory, the black line the mean trajectory for each type. Grey bars indicate day of flight. D) Proportion of travelers showing symptom trajectory types 1-4. E) Comparison of individual mean symptom scores per symptom domain before (7d) and after the flight (4d). F) Output from repeated-measures logistic regression predicting post-flight status from daily scores of each symptom domain. Odds ratios and their 95% CIs are indicated. Asterisks indicate p-values from Wilcoxon signed rank tests (B,E) or t-tests as per Tab. S1 for the predictors in F. *, p < 0.05; **, p < 0.01; ***, p < 0.001. All boxplots are Tukey boxplots.

Importantly, however, there was considerable variability in symptom load and trajectories between individuals. Mean symptom scores at baseline varied almost as widely as those post-flight (Fig. 1A,B) - although participants considered themselves generally as healthy (Tab. 1). In terms of symptom dynamics, we identified 4 types of trajectories (Fig. 1C): 2 types with an increase in symptom load post-flight (Type 1 and 2) and 2 without (Type 3 and 4), splitting the sample approximately in two (54% versus 46% of participants, Fig. 1D). The increase in jetlag symptoms for Type 1 was intense but short lived, while that of Type 2 persisted over the entire observation period post-flight. The few participants with Type 3 trajectories reported hardly any symptoms at any time, while the more numerous with Type 4 had a considerable symptom load throughout.

Were there particular symptoms driving these observed patterns? None of the 5 symptom domains (cognitive, affective, physical, sleep, and vegetative) or the underlying individual symptoms stood out as over-proportionally influencing the overall score trajectory, pre- and post-flight level differences or the 4 trajectory types. Although the absolute levels for each symptom and domain differed, the individual patterns were very similar to the total score patterns (Fig. 1E, S2A-D). Using logistic regression to identify the symptom domains most suited as markers for post-flight vs. pre-flight status, we found that increased sleep and vegetative symptoms were indicative of a post-flight status (if all other symptoms are held constant), while the isolated increase in cognitive symptoms was actually predictive of a *pre-*flight status (Fig. 1F, Tab. S1) likely due to the high prevalence of cognitive symptoms already pre-flight (Fig. 1E, S2A-D).

In summary, while jetlag syndrome can be described as the malaise following trans-meridian jet travel, it constitutes a complex phenomenon that cannot be sufficiently understood without considering individual pre-flight state and post-flight symptom dynamics.

### Quantifying jetlag based on intra-individual symptom scores: Intense jetlag differs from persistent jetlag

Aiming for a quantitative assessment of jetlag syndrome that accounts for different trajectories and baseline dynamics, we devised three simple metrics from the symptom time series: jetlag *peak intensity* and *mean intensity* (Fig. 2A), both baseline-corrected by subtracting individual mean/median baseline scores, and the *pivot* of jetlag, a center-of-mass measure for jetlag dynamic and duration (Fig. 2B). To gauge their suitability and behavior, we compared these metrics with each other as well as with 2 “traditional” jetlag measures that emulate previous jetlag quantification strategies and thus do not employ baseline correction and are based on standard single-question ratings of subjectively experienced jetlag (“How much were you impaired by jetlag symptoms?”). From a daily such question, we used the post-flight maximum as the *traditional peak metric* of jetlag, and from a retrospective such question applied to 43% of the sample, we gleaned the *traditional retrospective metric*.

**Figure 2.**
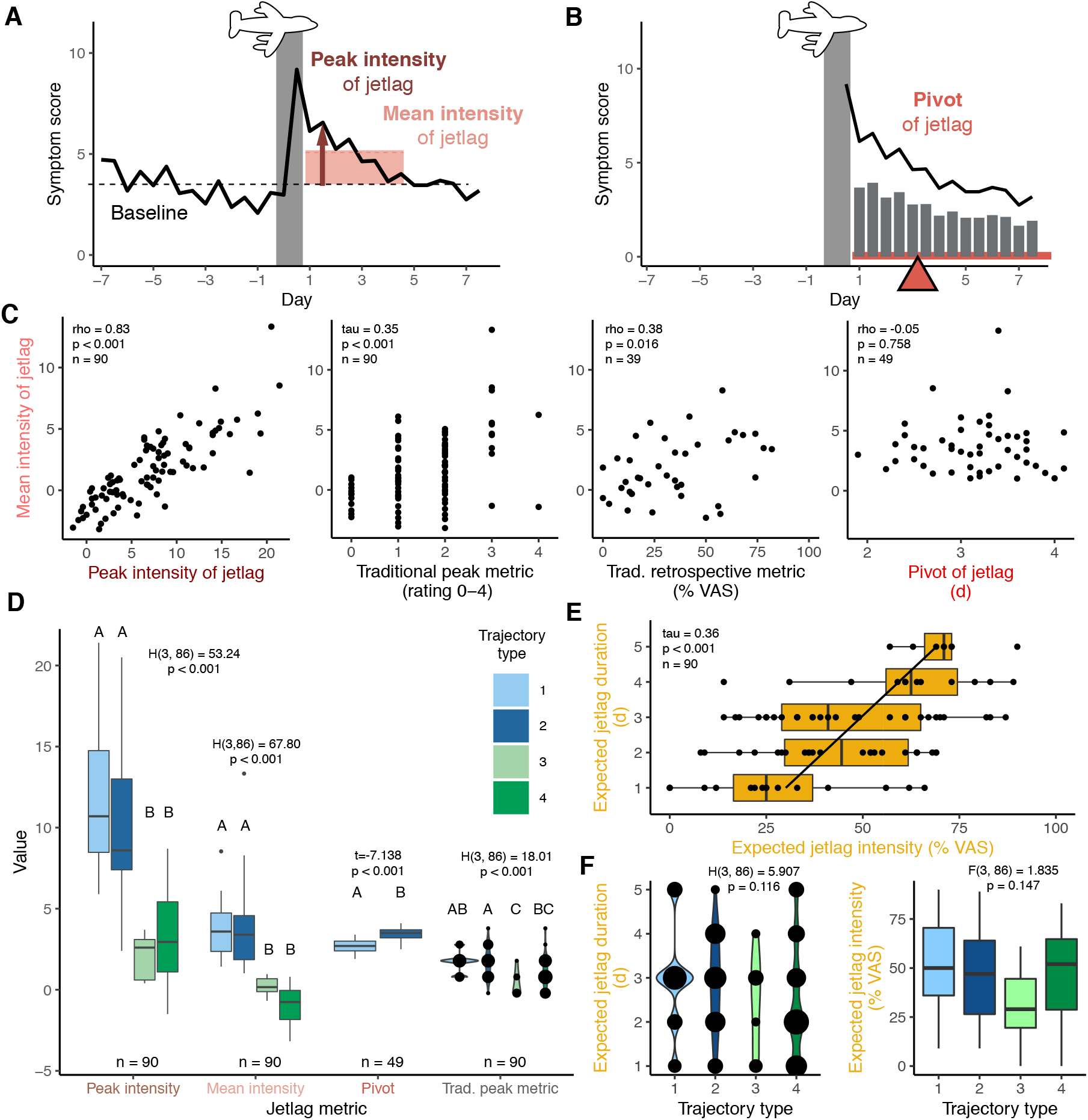
Jetlag quantification based on symptom trajectories. A,B) Schematics illustrating the newly devised metrics for jetlag quantification from symptom scores. Although the population mean (black, from Fig. 1A) is shown for illustrative purposes, all measures were calculated strictly on the individual level. A) Peak intensity of jetlag (dark red arrow) was defined as the *maximum* individual increase from individual median baseline (dashed line) *within* the first 4 days post-flight, mean intensity of jetlag (light red square) as the individual *mean* increase from individual mean baseline *across* the first 4 days post-flight. B) Pivot of jetlag (red triangle) as a measure of jetlag duration was defined as the center of mass of individual symptom scores post-flight. The pivot balances the weights of the symptom scores (grey bars; 50% of true weight for ease of visualization) as on a seesaw (red bar). C) Relationship among indicated jetlag metrics. Results of correlation analyses (Spearman rho, Kendall tau) are provided. D) Jetlag metrics segregated by symptom trajectory type as per Fig. 1C. Result of one-way ANOVA and Kruskal-Wallis test are provided. Letters indicate results of post-hoc pairwise comparisons, with data marked by different letters demonstrating significant differences after Bonferroni-adjustment. E) Relationship between the two jetlag expectation measures, expected intensity and expected duration. Regression line is solely for visualization. F) Expectation ratings segregated by symptom trajectory type. Left panel shows expected duration, right panel expected intensity. All boxplots are Tukey boxplots; in case of discontinuous distributions, these were replaced by bubble charts with bubble area encoding number of observations and visual support from violin plots. For better display, 3 extreme values of expected jetlag duration, which do not drive the results, were omitted from the graphs. Abbreviations: Trad., Traditional; VAS, visual analog scale.

Our analyses revealed expected and desired similarities and differences between the metrics. The *peak intensity* of jetlag, the *highest* increase in symptom load within the first 4 days post-flight, was at a median of 6.8 points (IQR: 2.9-10.2) over the entire sample, representing a ≥1-unit increase across ≤7 symptoms. The *mean intensity* of jetlag, the *mean* increase in symptom load over the first 4 days post-flight, was at a median of 1.3 points (−0.4-3.5), indicating an average ≥1-unit increase in ≤2 symptoms. These two intensity measures correlated both with each other as well as with the 2 traditional metrics (Fig. 2C, Fig. S3A), indicating their relatedness but also non-redundancy. Both peak and mean intensity also segregated with the 4 symptom trajectory types, distinguishing adequately between those with systematic symptom increases (Type 1,2) and those without (Type 3,4), whereas the similarly analyzed traditional peak metric frequently mis-quantified high general symptom variance as severe jetlag (Fig. 2D).

As a measure of jetlag duration, we opted for the center of mass of symptom scores post-flight, which we termed *pivot* of jetlag. As illustrated in Fig. 2B, the pivot marks the day post-flight at which the prior symptom load balances out the subsequent symptom load like on a seesaw. An earlier pivot corresponds to a fast jetlag dynamic with quickly subsiding symptoms, a later one to drawn-out jetlag symptomatology. Since assessing jetlag duration requires the presence of jetlag syndrome in the first place, we classified travelers into jetlagged/not-jetlagged based on a heuristic, intra-individual cut-off of jetlag mean intensity ≥1. Based on this, jetlag prevalence was 54% in our sample (Fig. S3B), a value similar to the 60% determined by Becker et al. with the same questionnaire using their own *inter*-individual split criterion ^30,35^ and also close to the estimate by Arendt et al. ^20^. In the 54% deemed jetlagged in our sample, the pivot was located at a median of 3.2 days (IQR: 2.8-3.5) and distinguished indeed between those with short (Type 1) and drawn out (Type 2) symptoms (Fig. 2D). Interestingly, the pivot of jetlag was not correlated to any of the jetlag intensity measures, neither the new baseline-corrected ones from detailed symptom scores nor the traditional ones without baseline-correction from single question ratings (Fig. 2C, Fig. S3A). This implies that jetlag intensity and duration may be independent aspects of jetlag, possibly reflecting differential regulation or modification by circadian re-entrainment.

Taken together, the above results demonstrate that finer jetlag quantification based on symptom scores is feasible and easy to implement with sufficient baseline information. Both peak and mean intensity quantify symptom increase post-flight, while the pivot reflects an intensity-independent duration of jetlag symptoms.

### Travelers expecting intense jetlag expect also long jetlag – independent of their symptom trajectories

What did travelers expect about their jetlag? Before their flights, participants expected to experience a jetlag intensity at 46% of the VAS scale (median; IQR: 28-65) and a median symptom duration of 3 days (IQR: 2-3; Tab. 1, Fig. 2E). Notably, expected jetlag intensity and duration were correlated (Fig. 2E), indicating that participants may have assumed a link between those two phenomena that we had not found in our quantification of the subsequently exhibited jetlag (Fig. 2F, Fig. S3). Moreover, travelers apparently did not anticipate their subsequent symptom trajectories since neither expected jetlag intensity nor duration differed between the trajectory types (Fig. 2F).

### Expectation is more predictive of jetlag severity than classic circadian parameters

In order to model the influence of jetlag expectation on jetlag syndrome severity, we performed multivariate linear regressions. As outcomes, we used the 3 new jetlag metrics (*peak intensity, mean intensity, pivot*) as well as the *traditional peak metric*. As independent variables (predictors), we included the general characteristics *age, gender* and *health*, the circadian characteristics *number of time zones crossed, direction of travel* (east/west) and *chronotype* (corrected midsleep times on free days, MSF_sc_), as well as one of the two expectation characteristics, either *expected jetlag duration* or *expected jetlag intensity* (Fig. 3A). We also included an interaction term between *chronotype* and *direction* to model chronotype-specific direction effects. This constellation provided us with a statistical power of 91%, at the Bonferroni-adjusted alpha-level of 0.00625 for 8 planned models (4 outcomes x 2 expectations), to detect a large effect of all predictors combined, an effect size we certainly expected given the many classic determinants included in the analysis.

**Figure 3.**
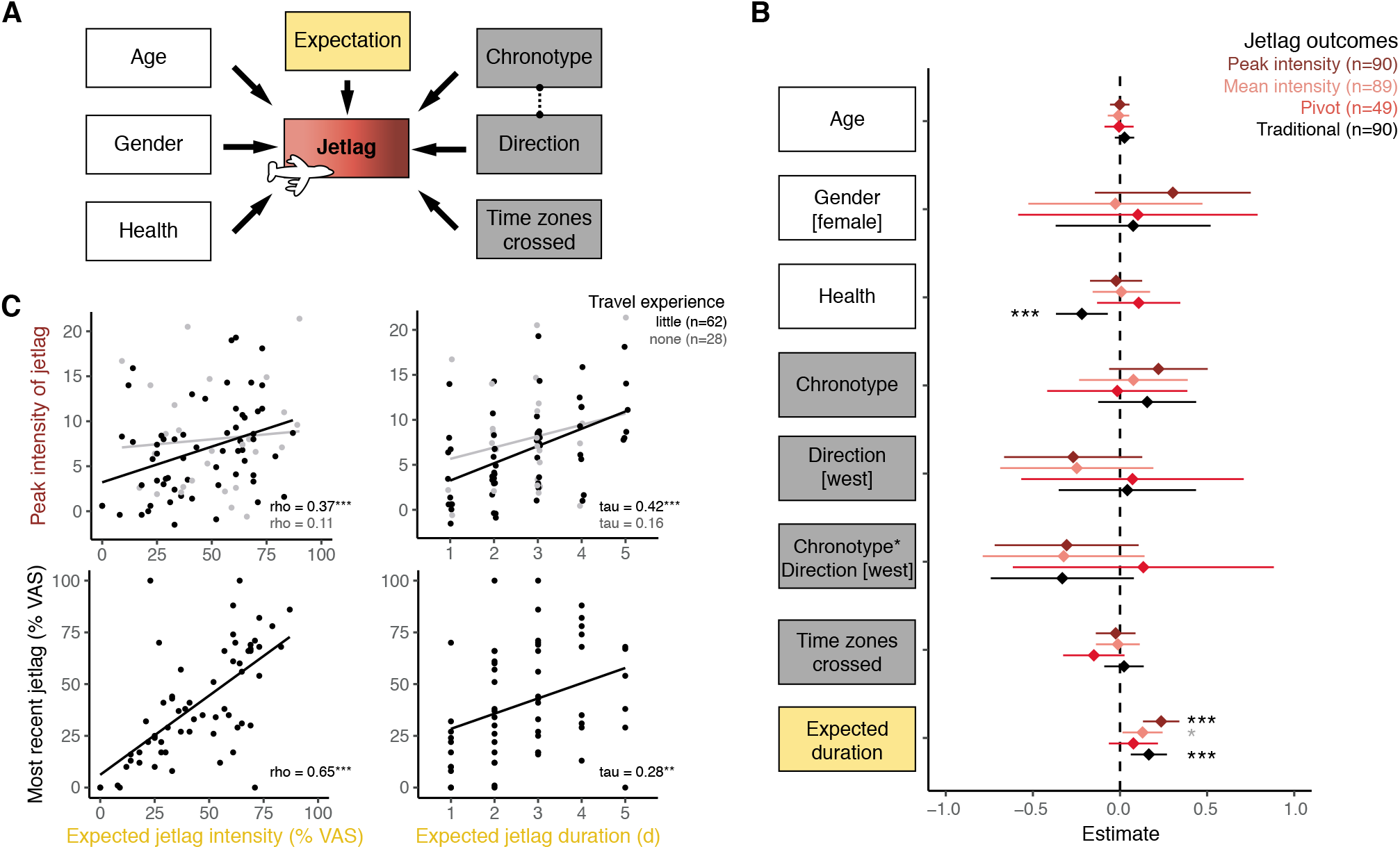
Jetlag syndrome prediction via regression models. A) Schematic of the regression analyses modelling the influence of individual predictors on various measures of jetlag symptom intensity and duration. As predictors (independent variables), we included general characteristics (white), circadian characteristics (grey), and expectation (yellow). B) Forest plot showing the regression estimate (± 95% CI) for each predictor (y-axis) and for the 4 main jetlag outcomes (color-code). Estimates are displayed for the models with expected jetlag duration (Table S2) and standardized, untransformed outcomes to enable direct comparison between outcomes, while predictors are unstandardized, i.e. coded in their respective unit of measurement (Age: y; Health: 1/10 VAS; Chronotype: h from mean chronotype i.e. MSFsc; Expectation: days) to better gauge actual effect size. The interpretation is as follows: For every 1-unit change in the predictor, the jetlag outcome changes by the indicated number of standard deviations. Note also that chronotype represents the chronotype effect on eastward travel because of the included interaction term chronotype*direction, which encodes the westward chronotype effect. C) Relationship between jetlag expectation, quantified jetlag severity on this journey and reported jetlag severity after the last long-haul journey for participants with little (black) and no (grey) long-haul travel experience (flights >3 time zones). Correlation results (Spearman rho or Kendall tau) are indicated, regression lines are solely for visualization purposes. For better display, 3 extreme values of expected jetlag duration, which do not drive the results, were omitted from the graphs. Abbreviations: VAS, visual analog scale; MSF_sc_, midsleep on free days sleep corrected. *, p < 0.05; **, p < 0.01; ***, p < 0.001; grey marks values above the adjusted alpha level of 0.00625.

Much to our surprise then, none of our 8 regression models demonstrated much explanatory power for any of the jetlag outcomes (Fig. 3B, Tab. S2-3). Only 3-28% of variance (R^2^) in jetlag syndrome severity was explained per model, with most predictors – including the circadian parameters – remaining far from statistical significance.

Notably, among the two predictors that showed at least some systematic explanatory power was health and expectation (Fig. 3B, Tab. S2-3). While health status predicted only the coarser traditional peak metric, expected jetlag duration significantly predicted both the traditional peak metric as well as the peak intensity metric – also after stringent adjustment for multiple testing (alpha = 0.00625). Namely, for each increase in expected duration by 1 day, the model showed (i) that the traditional metric increased by 0.2 on its 5-point scale (b=0.16; p=0.002,), and (ii) that jetlag peak intensity increased by 1.3 units, i.e. one symptom by one quarter (b=1.27, p<0.001, Tab. S2). Over the core range for the duration-expectation from 1 to 5 days, this effect would correspond to approximately a doubling of the peak intensity from a median of 6.8 units to >13 units. For the other two jetlag outcomes, which are not concerned with symptom peaks, there was, however, no direct evidence for an effect of expected jetlag duration. The effect on jetlag mean intensity did not pass adjustment for multiple testing (b=0.34, p=0.028), while there was no effect at all detectable for the pivot of jetlag, the actual metric for jetlag duration (b=0.04, p=0.277, Tab. S2). Notably, regression models including expected jetlag *intensity* instead of *duration* yielded comparable signals, albeit with weaker effects for expectation that did not pass the adjustment for multiple testing (Tab. S3). Still, expectation effects were the only ones besides health status to pass even the unadjusted alpha-level of 0.05. In summary, expected jetlag duration was systematically linked in our sample with jetlag intensity but not duration, whereas effects for expected jetlag intensity pointed in the same direction but did not reach statistical significance after adjustments.

### Predictive merit of expectation might be linked with previous experience

Was the effect of expectation on jetlag intensity based on participants’ prior jetlag experience? Our exploratory analyses in this direction certainly indicate that past experience may have played a role. Namely, both expected jetlag intensity and duration were correlated with jetlag intensity only in travelers that had some prior long-haul travel experience (n=62) but not in those that were completely naive in this regard (n=28; Fig 3C). Furthermore, in those with experience, there was a strong correlation between their most recently experienced jetlag severity and their jetlag expectation in this study (Fig. 3C). This hints that jetlag may either manifest itself to an extent expected from prior experience (placebo/nocebo effects) or that jetlag severity may be somewhat trait-like and individual characteristics may be more deterministic than travel parameters.

### Circadian parameters show predictive value for sleep parameters from diary but not for jetlag outcomes

Faced with the unanticipated poor predictive power of our models and particularly that of the circadian parameters, we sought to rule out an unsuitable jetlag outcome as the core reason. We thus performed further, exploratory regression analyses on alternative jetlag outcomes. We used (i) a dichotomous jetlagged/not-jetlagged classification (Fig. S3B), (ii) quantified jetlag based on individual symptom domains including those that had demonstrated predictive power for a post-flight status (sleep and vegetative symptoms, Fig. 1F) as well as (iii) enlisted principal component analysis for a data-driven re-combination of symptoms into new domains (cognitive, sleep, vegetative symptoms; Fig. S4A-D) to quantify jetlag on new individual clusters.

Regression analyses on these 9 alternative jetlag outcomes did not reveal broadly systematic effects on jetlag either (Tab. S4). At the adjusted alpha-level of 0.0056, the interaction for chronotype and travel direction showed a predictive effect for two of the outcomes, the symptoms from the original physical domain as well as the new vegetative symptom cluster from PCA. Here, westward travel led to smaller symptom increases the later an individual’s chronotype (b= −0.45, p=0.006 and 0.008). Also, expected jetlag duration had a predictive effect on the physical domain outcome, with each additional expected day explaining a 0.13-unit increase in symptom load (b=0.13, p=0.001). Thus, the detected effects were in expected directions but far and few between.

When looking at quantitative sleep outcomes from daily sleep diaries, however, the same regressions yielded more convincing effects in line with circadian sleep biology ^36^ (Tab. S5). Although number of time zones crossed still showed no predictive value, travel direction emerged as a good predictor for changes in sleep latency, sleep efficiency and midsleep post-flight, partially paired with chronotype. Namely, in westward travel, sleep latency increased less than in eastward travel (9 min smaller increase, b=-9.24, p<0.001), sleep efficiency was less reduced (3% smaller reduction, b=3.21, p=0.009), and midsleep times (in respective local time) were advanced by 1.3 h (b=-1.32, p<0.001), in particular for late chronotypes (0.6h more per 1h later chronotype, b= −0.61, p=0.008). On easterly journeys, later chronotype was almost predictive of a greater reduction in sleep efficiency (2% greater reduction, b=-2.16, p=0.018), just missing the adjusted p-value cutoff of 0.0125 for 4 sleep models. Notably, expectation demonstrated no predictive value for any of these sleep outcomes (Tab. S5).

## Discussion

Jetlag is probably the most palpable form of circadian misalignment. For some only a nuisance on their leisure trip, jetlag can become a serious hazard for persons in performance- or safety-critical tasks ^37-40^; under chronic exposure such as experienced by flight crew, jetlag might have even direct long-term health consequences ^41-44^. We thus require a solid understanding of the processes involved in jetlag and the individual factors influencing susceptibility and symptom severity. Our aim here was to investigate whether and to what extent jetlag symptomatology may be influenced by prior expectation - to inform both mechanistic understanding and treatment improvements in jetlag disorder.

In our sample, only 54% of participants experienced a post-flight malaise despite everyone crossing >3 time zones. This proportion is in line with the three previous reports on jetlag prevalence that we know of ^20,30,35^ but is in stark contrast to the apparent wide awareness in the general population about the syndrome and also the expectation in our sample, where over 75% anticipated a jetlag severity of more than 25% of the scale. This strikingly low prevalence, although no large-scale data exist, poses interesting questions in terms of inter-individual differences in jetlag disorder.

High variability in symptom load and trajectories was definitely evident in our sample. Based on the 4 symptom trajectory types that we identified, we developed 3 metrics to adequately quantify the extent of jetlag symptoms in light of this inter-individual variability: jetlag peak intensity, mean intensity and pivot of jetlag. Although these metrics still require external validation and are likely far from perfect, our plausibility checks in relation to other measures indicate that they are adequate representations of the symptom trajectories observed and are likely more suited to a fine-grained jetlag quantification than previous coarse measures.

Much to our surprise, the classic circadian factors number of time zones crossed or travel direction did not show any predictive power for the individual differences in jetlag symptom severity; the only factor explaining at least some of the variability in jetlag syndrome was expectation. Health status emerged also as a significant predictor but only for the baseline-uncorrected traditional peak metric, which is directly influenced by absolute symptom load, for which a link to health status is self-evident. The effect of expectation on jetlag symptoms was in the anticipated direction: the more severe participants expected their jetlag to be, the more severe the jetlag outcome was. Since jetlag outcomes were quantified based on subjective symptom reports, we cannot conclude at this point whether subjects who expected more jetlag were just more aware of their symptoms leading to the subjective experience of a higher or longer symptom load or whether symptoms were indeed more pronounced.

Interestingly, participants did not distinguish clearly between expected jetlag symptom intensity and duration. Both expectation measures were correlated, and expected duration predicted jetlag intensity not jetlag duration. The expected intensity showed a similar pattern albeit below the adjusted alpha-level for statistical significance. Furthermore, previous jetlag experience seemed to play a role in jetlag expectation and actual jetlag severity. This leaves room for various speculations on the driving mechanism behind these associations. Is jetlag trait-like in that individual characteristics are more important for jetlag severity (or jetlag symptom perception) than external parameters, so that past experience is a good indicator for future experience no matter how many time zones were crossed? Or does past experience determine future experience through psychological mechanisms?

Research on placebo effects has clearly shown that prior experience shapes expectations, and expectations, in turn, shape symptom severity ^45,46^. For example, the expectation of nausea prior to chemotherapy was shown to be modulated by prior nausea experience, while the severity of chemotherapy-induced nausea was predicted by nausea expectation ^47^. Our findings therefore have also clinical implications from the perspective of placebo research. First, expectations are not trait-like but can be targeted by psychological interventions. Psychological interventions could thus be useful to reduce jetlag syndrome. For example, the optimization of expectations in heart patients prior to surgery by a short psychological intervention improved the long-term outcome of surgery ^48^. Second, a recent study indicates that expectations and hence treatment outcome can be optimized by administering placebos ^49^. Especially open-label placebos that are delivered honestly, i.e., without deception ^50^, may be a promising approach to target the symptoms of jetlag disorder in the future.

A major question raised by our results is why were we unable to detect strong and consistent effects of classic circadian parameters on jetlag symptom severity? All regression coefficients pointed in the expected direction but the inter-individual differences were much too large for a systematic effect to emerge (Fig. 3B). The fact that we identified circadian effects on the sleep parameters from diaries – although certainly not broad or large effects – suggests that our sample most definitely experienced circadian misalignment as would be expected from basic circadian principles. There are several possible explanations why we might have nonetheless failed to detect impacts on the jetlag outcomes: (i) Our jetlag measures might not have captured the phenomenon appropriately despite our careful design. (ii) We might have omitted an important negative confounder (variable obscuring existing relationships because it is causally linked with both predictor(s) of interest and the outcome). (iii) The included predictors have indeed no substantial influence on jetlag severity.

i. Jetlag outcomes: In subjective symptom reports, there is an inherent uncertainty if reported symptoms match objective symptom occurrence or strength, as participants may under- or overestimate their symptoms (e.g. in fatigue/sleepiness studies ^51^). Therefore, even our carefully designed jetlag metrics as well as our additional exploratory outcomes (ranging from coarse classifications to detailed PCA) might not have overcome this bias so that we may have captured more the psychological than the physiological aspects of jetlag syndrome. It has to be noted, though, that our metrics were based on the integration of detailed longitudinal twice-daily symptom reports rather than a mere retrospective jetlag assessment using a single question as is done to capture the construct of “subjective jetlag” ^20-22,24,25,52^.
ii. Confounding: One might also argue that we did not include detailed information on light exposure in our models as it was not specifically collected in our study. Light as the main zeitgeber in humans is, of course, of paramount importance in re-entrainment, and thus would be expected to have a good predictive value at least for jetlag duration. However, to a great extent, this is exactly what the factors number of time zones crossed as well as travel direction encode: the experienced shifts in the natural light-dark cycle (photoperiodic changes were small in our cohort; Fig. S1C-D). Since the majority of participants showed almost instantaneous behavioral adjustment to the new time zone (as judged by midsleep time, Fig. S4E), number of time zones and direction should be a reasonable approximation of the actual light-dark-cycle shift experienced. Even under poor approximation, additional light information might increase the predictive value of the entire model but its absence is unlikely to obscure effects of the other predictors included.
iii. No substantial effect: If the answer for our difficulties in modelling jetlag severity is not inappropriate outcome measures or a missing confounder, it may well be the complexity of the relationship between circadian mis-/re-alignment and ensuing symptoms and their dynamics, i.e. jetlag syndrome. The focus on circadian re-entrainment in jetlag research has inherently assumed that the extent and rate of re-entrainment strongly determines the intensity and/or duration of manifested symptoms. But this may not be the case – as was already pointed out by Atkinson et al ^5^ but apparently not yet taken widely onboard. Importantly, we are also not the first to fail in modelling jetlag symptom severity based on classic circadian predictors: In a sample of 53 travelers, Becker et al. found no relationship between daily jetlag symptom scores and direction of travel, number of time zones crossed and chronotype using multi-factor MANOVA ^30^. Even in certain re-entrainment scenarios, some data even point towards limited predictive value of the classic parameters because of inter-individual variability in re-entrainment also in inbred animal models under controlled laboratory conditions ^19,53^ or mathematical models suggesting complex interactions between circadian parameters, which can lead to faster re-entrainment after more and not less time zones crossed ^54^. If our findings can be further replicated in other samples, this calls into question the current guidelines on the determinants of jetlag syndrome.

## Conclusion

In conclusion, our study raises many questions and highlights the need for several future lines of jetlag research. We require better quantifications of jetlag severity on a symptom level (i.e. jetlag syndrome), to which we contributed here with new measures for jetlag intensity and duration. We need to consider psychological mechanisms such as expectation effects directly in our studies, not just to compare a particular treatment against a placebo intervention but also to actively leverage placebo effects and reduce nocebo effects for better outcomes. Finally, we ought to differentiate better between the different jetlag constructs, i.e. change in Zeitgeber timing, resulting circadian mis-/re-alignment, and manifested downstream symptomatology. Investigations into the link between the latter two, a link likely complex and non-linear, should become a priority to enable better translation of circadian basic science – in jetlag and beyond.

## Acknowledgements

We thank the participants for their contribution, Prof. H Herzel for constructive feedback on preliminary results and suggesting the center of mass as duration measure and W. Schwartz for critical discussion.

## Author Contributions

Study conception and design: ECW, KM

Creation and implementation of the questionnaires: KM, SD, ECW

Acquisition of data: MU

Data Curation: MU

Analysis and interpretation of data: MU, ECW, KM, DF

Funding Acquisition: ECW

Writing:

-Drafting of Manuscript: MU, ECW

-Reviewing & Editing: ECW, KM, MU, DF, SD

Supervision: ECW, KM

Visualization: MU, ECW

## Methods

### Sample Details

#### Participants

##### Recruitment

Participants planning to fly over more than 3 time zones were recruited through convenience sampling via flyers, university webpages and mailing lists. Inclusion criteria were: ≥18 years old, no psychiatric disorder or any disorder affecting sleep, currently neither pregnant nor breastfeeding, no shiftwork within the 8 weeks preceding travel, no flights over >3 time zones within the year before the study and ≤4 times during the last 10 years (subsequently relaxed to ≤5 times), plan to stay at destination for ≥7 days without crossing further time zones and the aim to follow a “normal” day-night-rhythm at travel destination (i.e. no excessive partying). Participants received financial compensation for partaking in the study.

##### Sample

108 participants provided written informed consent confirming to fulfil the inclusion criteria, of which 18 participants were later excluded from the analyses due to the following reasons: 12 provided not enough data points (completed <27 of the 31 questionnaires); 4 stated a travel history of >5 flights over >3 time zones during the last 10 years in the baseline questionnaire; 1 protocolled only their return journey after a stay abroad and could therefore not be assumed to be free of jetlag at baseline; 1 had a 2-day layover between origin and destination.

These exclusions resulted in a final sample size of 90 participants. While most participants were the only one in their travel group enrolled in the study, in 12 instances 2 participants traveled together, i.e. 24 individuals in total. We specifically sought participants with limited trans-meridian-travel experience in order to minimize the impact of previous jetlag experience from similar journeys on jetlag expectation. This was important to interpret the directionality of any associations between expectation and jetlag severity.

Basic population characteristics as well as travel experience are provided in Table 1. The majority of our participants were university students (n=68), only 15 were employed, only 3 had children (aged 3-5 years). Five participants were smokers, 51 stated to work out/do sports on a regular basis, quality of life was at a median of 85% VAS (range: 50-100). 20 of the 58 women used oral contraceptives, 11 participants took vitamin supplements, 4 thyroid hormones, and 3 medication against asthma.

##### Ethics

The study was performed in accordance with the Declaration of Helsinki and was approved by the ethics committee of the Medical Faculty, Ludwig Maximilian University (17-350).

#### Travel Details

Flights originated from European airports (mainly Munich) and were headed to various destinations around the world a median of 6 time zones away and approximately equal proportions of easterly and westerly flight directions (Fig. S1A, Tab. 1). Flights occurred between January 2018 and November 2018, clustering during university breaks in spring and early fall (Fig. S1C).

Since photoperiodic change has been postulated as a potential factor in jetlag re-entrainment based on a mathematical model ^54^, Figures S1C-D provide a graphical display of the photoperiod changes experienced by our sample. Since most flights occurred close to the equinoxes and involved mainly small changes in latitude, there was little change in photoperiod for most travelers, hence we decided against including photoperiodic change as a factor in our regression models (see below).

Participants were not instructed on causes, symptoms or countermeasures of/for jetlag by the study team. To our knowledge, participants did not intentionally apply any measures to alleviate jetlag symptoms such as oral melatonin or light therapy.

### Method Details

#### Study Design

This study was designed as an observational study with prospective monitoring of jetlag symptoms over 2 weeks. Upon enrollment, all participants filled in an entry questionnaire assessing basic sociodemographic characteristics, medication, sleep and chronotype as well as travel details, travel experience and jetlag expectation. Seven days before their respective flight, participants started recording their jetlag symptoms twice daily and continued this until 7 days post-flight.

#### Questionnaires

##### General

Questionnaires were administered in German and online using REDCap electronic data capture tools hosted at the LMU Munich ^55,56^. Immediately after enlisting in the study, participants received a link to the entry questionnaire. For the twice-daily jetlag questionnaires, the system was set up such that questionnaires became accessible only on the appropriate day. Since this mechanism did not account for shifts in time zone, some participants gained access to their questionnaires slightly earlier than appropriate for their new destination leading to some participants with data only until the 6^th^ day post-flight (see missingness matrix Fig. S1B). Submission of each questionnaire was time-stamped and was only possible once all questions had been completed. If participants anticipated an unstable internet connection at their travel destination, we provided hard copies of the questionnaires and enabled retrospective population of the online forms based on the manual notes (n≈24 participants).

##### Entry Questionnaire

The entry questionnaire was a combination of several individual questionnaires and questionnaire items including (i) demographic assessment (age, gender, height, weight, family status, number and age of children, highest educational qualification, employment status, habits concerning smoking, alcohol and sports, medication, general health status and perceived quality of life), (ii) flight details (origin, destination, timing of take-off and landing, direction of travel, time difference between origin and destination, (iii) flight and jetlag history (whether this was the participant’s first flight over >3 time zones, how many of such flights the participant experienced within the last 10 years, the extent of impairment by jetlag symptoms following the last flight over >3 time zones), (iv) the Munich ChronoType Questionnaire (MCTQ ^33,34^) for baseline circadian assessment and chronotype determination via the midsleep time on free days corrected for oversleep (MSF_sc_), and (v) jetlag expectation assessment.

##### Jetlag Expectation

Jetlag expectation was assessed via 2 central questions in the entry questionnaire. These questions were modelled after previous expectation questions that commonly assess the expected intensity and duration of an effect ^e.g. 57^: (i) “How strongly do you expect to get impaired by jetlag symptoms with your upcoming flight?” (expected jetlag intensity) with answers on a visual analogue scale (VAS) given their ease of use and avoidance of imprecise terminology ^e.g. 58^. (ii) “After how many days do you think your jetlag symptoms will have subsided?” (expected jetlag duration). The answers to the second questions contained 3 outliers outside the range of the other points between 1-5 days (2x 10 days, 1x 14 days; >3 standard deviations from the mean). Since these outliers nonetheless represent plausible values that can be seen as part of the right-tail of the distribution, we included the outliers in our analyses but ran sensitivity analyses with those excluded and found essentially equivalent results. For a useful graphical display of expected duration, we removed the outliers from the graphs.

##### Charité Jetlag Scale

For jetlag monitoring, we used the Charité Jetlag Scale (CJS) ^30^, one of a few comprehensive jetlag questionnaires that assess the strength of a range of common jetlag symptoms over time. The CJS is a validated German version of the Columbia Jetlag Scale ^31^ including several small modifications based on the Liverpool Jetlag Scale ^32^.

The CJS was administered twice a day, once in the morning and once in the evening ^59^, for 7 days before the flight (days −7 to −1), on the flight day (day 0) and 7 days after the flight (days 1-7), resulting in 30 CJS that could be filled in. Morning questionnaires were marked with the full day number (e.g. day 1), evening questionnaires were designated via “.5” (e.g. day 1.5). To avoid an undue influence of the travel conditions on jetlag quantification, questionnaires pertaining to the flight day (day 0 and 0.5) were excluded from all analyses.

From a total of 19 symptoms clustered into 5 symptom domains (sleep, mental, physical, vegetative, and cognitive), each instance of the CJS assesses the severity of 15 of these, with the morning questionnaire containing 4 sleep-related items (3 items from the sleep domain and nocturia from the vegetative domain) that are replaced by 4 cognitive items in the evening questionnaire. Each item is rated on a scale from 0-4 (“not at all”-”very strong”); the total symptom score is the sum of all 15 symptom ratings within a questionnaire. The total symptom score thus ranges between 0 (no symptoms observed, score 0) and the maximum value of 60 (all symptoms observed, very strong, score 4). The internal consistency across items of the CJS was shown to be high ^30,35^, however, its individual items have not been validated against objective measures for each symptom except for the sleep items ^35^.

Morning and evening scores were kept separate and not aggregated into a daily score, as to retain the half-day resolution and avoid artificial designation of day boundaries, except for principal component analysis (see below).

##### Sleep Diary

In addition to the daily CJS, participants received a daily sleep diary with their morning questionnaire. This was a modification of the published CJS-associated sleep diary ^30^ altered to incorporate important concepts and variables from the MCTQ ^33,34^. Namely, the sleep diary consisted of questions on the time when participants went to bed (bedtime), when they were ready to fall asleep (lights off), how long it took them to fall asleep (sleep latency), the time when they awoke at the end of the night’s rest (wake-up time), when they got out of bed the last time before starting the day (get-up time), whether they were actively woken or not, how many times they woke during the night (awakenings), how many hours they actually slept during the night (sleep duration).

For analysis of parameters from the sleep diary, the variables sleep duration, awakenings, and sleep latency were directly taken from the questionnaire. Time of sleep onset was determined by adding the sleep latency to the lights-off time. Sleep efficiency was the ratio of sleep duration and the duration between lights-off and final wake-up, i.e. reflecting the proportion of time asleep while trying to sleep. The time of midsleep was the halfway point between sleep onset and wake-up time.

##### Correct assignment of day type

To ensure correct and consistent questionnaire assignment to day-number pre/post-flight, we manually crosschecked the automatic timestamp and the participant-provided current date, number of days to/passed since the flight, travel details and time differences, from which we generated the most plausible “correct” day assignment variable. Although this meant omitting some questionnaires that were filled in during a stopover leading to some participants having 1-2 fewer data points (Fig. S1B), this ensured that day numbering was consistent across all participants, with the designated Day 1 always the first day after travel.

### Quantification and Statistical Analysis

Data were processed, visualized and analyzed in R ^60^, and graphs plotted with the R package “ggplot2” ^61^. Other R-packages used for analyses and display were: car ^62^, circular ^63^, dunn.test ^64^, eulerr ^65^, Hmisc ^66^, lme4 ^67^, lmerTest ^68^, lubridate ^69^, maps ^70^, MESS ^71^, purr ^72^, pwr ^73^, reshape2 ^74^, sjlabelled ^75^, sjmisc ^76^, sjPlot ^77^, tidyverse ^78^.

Measures of center, dispersion and uncertainty are specified in the respective figures or their legends. All presented boxplots are Tukey boxplots, for which the box ranges from the 25^th^ to the 75^th^ percentile, the line marks the median, and the whiskers encompass all data points within 1.5 times the interquartile range while data points outside the whisker range were plotted as outliers. All hypothesis testing was performed two-sided with an alpha level of 0.05 unless Bonferroni-adjustment is indicated (see below).

#### Jetlag Quantification

The severity of jetlag, i.e. its intensity and duration, was quantified via 3 newly devised metrics (peak intensity, mean intensity and pivot) and 2 traditional metrics (traditional peak metric, traditional retrospective metric). To avoid an undue influence of the travel conditions on jetlag quantification, none of the measures took values of the flight-day into account (day 0 and day 0.5).

##### New jetlag metrics

The new jetlag metrics were calculated from total symptom scores from the CJS. For peak and mean intensity, each symptom score post-flight was baseline-corrected by subtracting the participants’ mean or median pre-flight score (mean for mean intensity, median for peak intensity; see below for rationale). *Mean intensity* was then defined as the *mean* baseline-corrected score over the first 4 days post-flight, *peak intensity* as the *maximum* baseline-corrected symptom score within 4 days post-flight as per the equations below. The 4 day-limit was chosen to cover the main period of symptom occurrence for most travel conditions ^4,59,79^. To account for the fact that morning and evening questionnaires contained 4 different symptoms, which resulted in systematically higher symptom scores in the evening as these symptoms generally received higher ratings (cf. Fig. 1A) ^59^, the baseline for peak intensity was determined using the median over all pre-flight ratings only from the same time of day (morning/evening) as the highest rating post-flight. The median was chosen over the more sensitive mean as it is less vulnerable to single outliers, which are more potent given the reduced number of baseline questionnaires due to daytime-selection.

The *pivot of jetlag*, as a measure of jetlag duration and dynamic, was calculated analogous to the center of mass. To this end, the symptom score for each day after the flight (the weight) was multiplied with its day number post-flight (the position). Those products were subsequently added up and divided by the sum across all post-flight symptom scores of that individual. The resulting number has the unit days and marks the point in time around which the symptom load is hinged, with a later pivot corresponding to more persistent symptoms. Importantly, the pivot of jetlag was only determined for participants with a jetlag mean intensity ≥1, since a prerequisite for a jetlag duration is a jetlag in the first place (see “Criteria for Jetlag Classification” below).

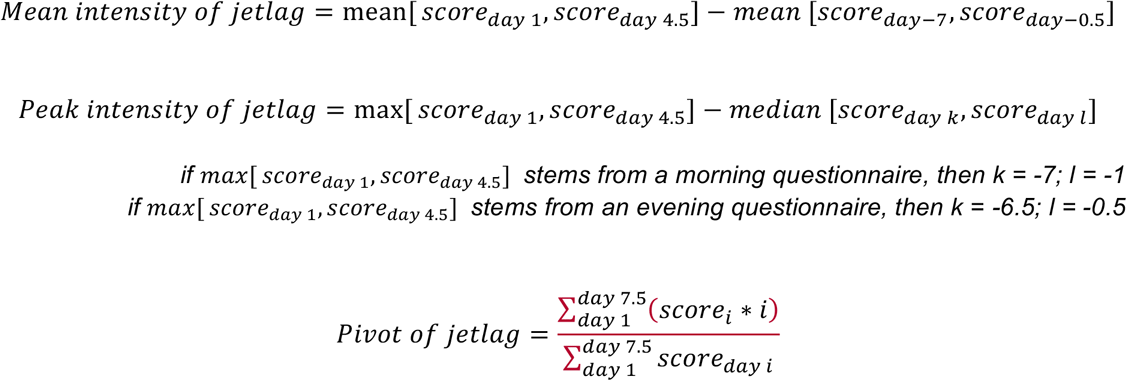

##### Traditional metrics

Each instance of the CJS contains a final question pertaining to the overall experienced impairment by the jetlag symptoms at that particular time point, rated on a scale 0-4 (“not at all” – “very strongly”). This is similar to other commonly-used coarse daily ratings of jetlag intensity ^9,28,80,81^. A single overall value of jetlag intensity was determined from these daily ratings analogous to the approach by Becker et al. for jetlag classification ^30^, taking simply the absolute maximum value across the post-flight period (days 1-7.5), which we loosely termed *traditional peak metric*.

For the last 43% of the sample (n=39), we added a summary question to the questionnaire on the final post-flight day. It asked for a retrospective rating of the overall experienced impairment by jetlag on a VAS. This question was added to enable comparisons of our new metrics with a second traditional metric, a retrospective subjective jetlag rating ^20-22^, which we loosely termed *traditional retrospective metric*.

##### Criteria for Jetlag Classification

For the binary distinction *jetlagged/not jetlagged*, we used 2 alternative criteria. The first was based on the mean intensity of jetlag, which we required to be ≥1 in order for an individual to be classified as jetlagged. This heuristic cut-off was based on the rationale that someone suffering from jetlag disorder should be expected to show at least a mean increase of 20% (1-unit on the 5-unit scale) in 1 out of 15 symptoms over the first 4 days post-flight. The alternative criterion was based on the traditional peak metric, which we required to be ≥2 (i.e. at least “modest” on the 0-4 rating scale) in order for an individual to be classified as jetlagged. This means the participant indicated at least once after the flight that they perceived at least a modest jetlag impairment.

Notably, there is a considerable 67%-overlap between travelers classified by either criterion (Fig. S3B). However, the traditional criterion marks also travelers with Type 3 and 4 trajectories as jetlagged, which do not show any systematic symptom increases, while the criterion based on the mean intensity by definition only marks Type 1 and 2 travelers as jetlagged (Fig. S3C).

##### Further Jetlag Quantification for Exploratory Analyses

For additional, exploratory regression analyses of other jetlag outcomes, we quantified jetlag from additional variables: (i) from the symptom score of each of the 5 CJS symptom domains (sum of all scores per domain), (ii) for each of the 3 principal components determined via principal component analysis (see below) as well as (iii) for the sleep variables nightly awakenings, sleep latency, sleep efficiency and midsleep time. To this end, we determined the mean intensity of each of these variables, namely the mean rating of the variable over the 4 days post-flight minus the mean rating of that variable over all days before the flight.

#### Jetlag Trajectory Types

Visual inspection of individual symptom trajectories revealed 4 different trajectory types in our sample. These were subsequently quantitatively assigned to each participant using the following rules: First, participants were divided into two groups, those with an increase in jetlag symptoms after the flight (mean intensity ≥1) and those without (i.e. jetlagged or not). Those participants with symptom increases were assigned to Type 1 if the mean symptom score of day 1-3 post-flight was at least twice the mean score of days 4-7. Otherwise the participant was assigned to Type 2. Those participants without an increase in their symptoms were either labeled as Type 4 if they had a symptom score that exceeded the arbitrary cutoff of 5 at least once within the entire observation period, or otherwise as Type 3.

#### Comparison of Means

Testing for differences in group means or medians between pre- and post-flight symptom scores and trajectory type-specific jetlag metrics was performed using the Wilcoxon signed rank test for paired samples, unpaired t-test, one-way analysis of variance (ANOVA) and Kruskal-Wallis test as indicated in the figures and their legends (after checks of the underlying assumptions). Statistically significant omnibus tests were followed by post-hoc pairwise-comparisons via t tests (ANOVA) or Dunn’s tests (Kruskal-Wallis).

#### Correlation Analyses

Associations between variables were assessed via Spearman rank order correlations including all pairwise complete observations. Rank order correlations were chosen to accommodate variables not normally distributed or at ordinal scale. In cases of many ties, which impairs ranking, we used Kendall’s tau instead.

#### Regression Analyses

##### Linear Regressions

Linear regressions were performed for continuous outcomes (jetlag metrics and sleep variables). The set of predictors was determined a priori based on theoretical considerations. The assumption of non-multicolinearity between predictors and/or outcomes was checked via correlation matrixes and scatterplots of all predictors and outcomes and was deemed fulfilled. This way, also non-linear relationships between any predictor and outcome variable was ruled out. Homoscedasticity was assessed by visual inspection of scatterplots as well as checking model residuals for normality via histograms and Shapiro-Wilk test. If a model’s residuals were not normally distributed, we removed any outliers. In the regressions for the following outcomes, exclusion of outliers was both necessary and sufficient: jetlag mean intensity (1 outlier removed), principal component 3 (1 outlier), delta sleep latency (5 outliers), and delta sleep efficiency (2 outliers). In the case of non-normally distributed residuals in the absence of outliers, as was the case for jetlag peak intensity, the outcome was log-transformed after adding a constant of 3 to raise all outcome values to ≥1: y=log_10_(x+3). Neither outlier exclusion nor log-transformation substantially altered model results.

Model structures, number of observations and key model parameters are all provided together with the estimates for standardized and unstandardized outcomes in Tables S2-5. For ease of interpretation, both the log-transformed and untransformed peak intensity models are listed, while Fig. 3B shows the estimates for untransformed peak intensity only.

##### Logistic Regressions

Logistic regressions were performed for the two dichotomous outcomes, (i) pre-flight/post-flight status and (ii) jetlagged/not jetlagged. Pre-flight/post-flight status prediction was performed on the symptom scores for each symptom domain (sum of all scores per domain) using the 5 domains as predictors to identify the symptom domains whose increase is most indicative of a post-flight status and thus potentially most indicative of jetlag disorder. For maximum information, individual scores for each questionnaire instance were entered into the model rather than pre-/post-flight aggregates, thus mixed model regressions were calculated with ID as random intercept (random effect) to account for the repeated-measures nature of the data. However, ID showed no random variance (singularity) and thus standard logistic regression would have yielded the same results. Prediction of jetlagged/not jetlagged was performed using standard logistic regression with the same set of predictors as the linear regressions above.

##### Adjustments for multiple testing

To reduce the family-wise error rate inflated by multiple regression models on similar outcomes, we performed Bonferroni-corrections based on the respective number of related models. The alpha-level for the 8 main models on 2×4 jetlag outcomes was thus 0.00625, for the 9 models on alternative jetlag outcomes 0.0056 and for the 4 sleep models 0.0125 as indicated in the text and respective regression tables (Tab. S2-5).

#### Principal Component Analysis

In search of a more sensitive jetlag outcome, we attempted to re-combine the 19 individual symptoms of the CJS into new, potentially more informative domains based on their covariance; in the original CJS they are assigned based on their physiological spectrum. We thus performed principal component analysis (PCA), entering the scores for each day (mean of morning and evening scores for symptoms rated both times and the actual value for symptoms rated only once a day) and person within the observation period as pseudo-independent measure (n=1295). PCA on repeated measures data is valid as data reduction technique but limits inferences on underlying clusters of symptoms ^82^. We confirmed the assumptions of sphericity, sufficient interactions between single symptoms and absence of multicollinearity via Barlett’s test, the Kaiser-Meyer-Olkin measure of sampling adequacy and the determinant of the underlying correlation matrix. Based on the symptoms’ eigenvalues, scree plot and overall fit of the model, 3 components were deemed the most suitable number of components. Since, in the context of jetlag, the 3 symptom clusters will hardly be independent from one another, oblique rotation was used. For scree plot and scatterplots of principal components, see Fig. S4A-D.

#### Power Analysis

A priori and post-hoc power analyses for the linear regressions were performed, using the R package pwr^73^. For a priori analyses, a regression model was simulated with 8 predictors and 90 observations, based on the final sample size and intended number of predictors. The alpha level was set to 0.00625 based on Bonferroni-correction for 8 planned models: 2 models for each of the 4 main jetlag outcomes, once with expected jetlag *duration* and once with expected jetlag *intensity* as predictor (see above). For small, medium and large effect sizes, the effect size f2 was set to 0.02, 0.15 and 0.35, respectively, yielding powers of 91%, 38% and 2.1%. We also performed post-hoc power analyses, keeping the parameters for the number of predictors and the alpha level but setting the power to 0.8, and f2 = R^2^/(1-R^2^), with R^2^ taken from the regression models. To reproduce our findings with a power of 80%, we would have needed the following number of participants: 72 for peak intensity, 183 for mean intensity, 193 for pivot, and 69 for the traditional peak metric (with expected duration as predictor).

## Resource availability

### Data and Code Availability

The data for all analyses and figures reported in this paper are available at [name of repository] [accession code/web link]. Further information and requests for resources should be directed to and will be fulfilled by the corresponding author, Eva Winnebeck.

