## Supplemental Material for "Jetlag Expectations, not Circadian Parameters, Predict Jetlag Symptom Severity in Travelers"

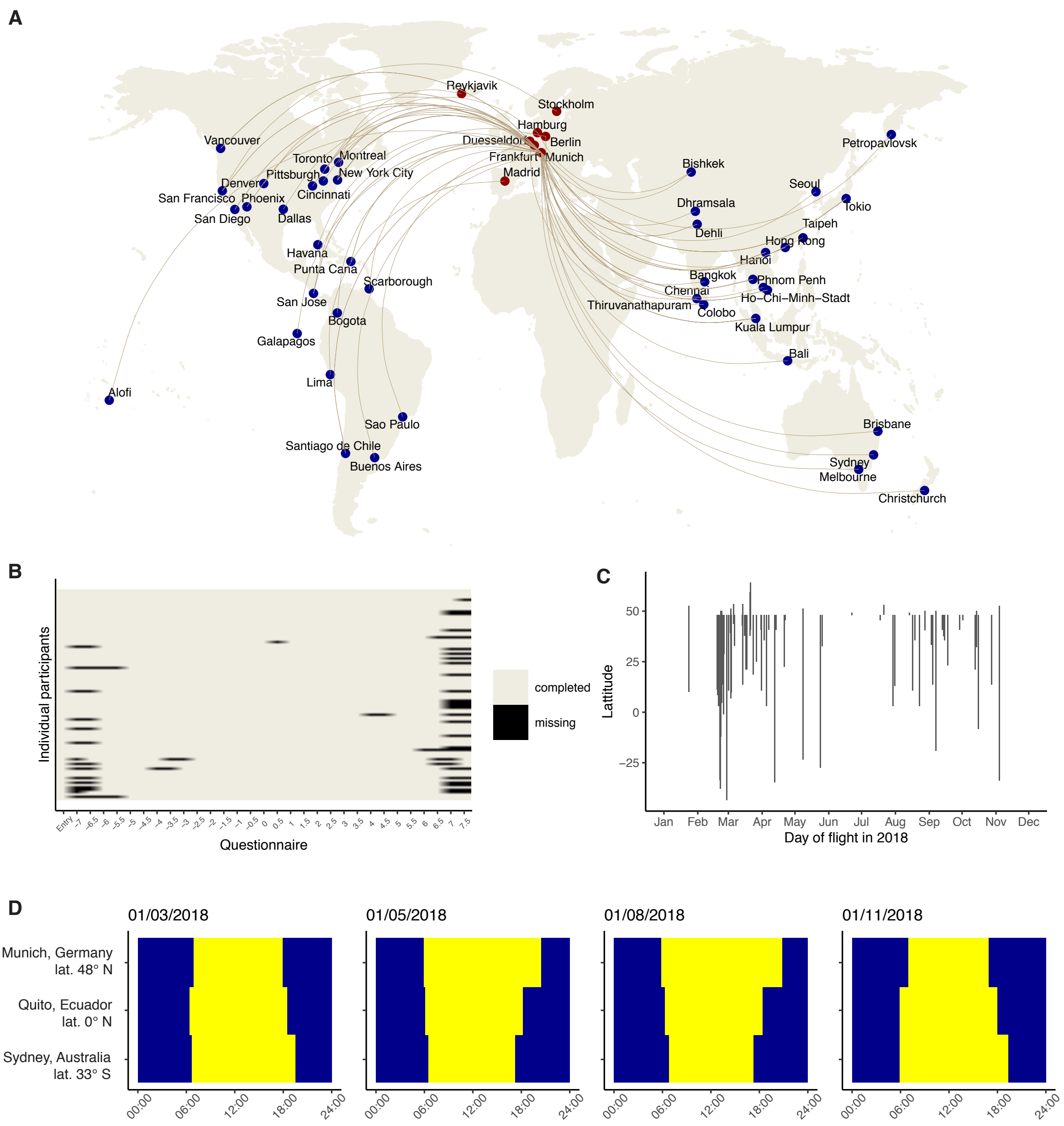


**Figure S1. Travel and data details. Related to Table 1.**
A) World map depicting travel routes of participants (n=90), with departures from central European airports (mainly Munich, red) and destinations around the globe (blue). B) Missingness map for all participants and questionnaire data points. Numbers on the x-axis indicate the day number in respect to the flight day (day 0). C) Time of year and changes in latitude for each travel route as indicator for experienced photoperiodic changes. Each line demarks one flight, spanning from the departure to the destination latitude. D) Photoperiods for Munich (the main travel origin), for Quito (representing equatorial regions) and Sydney (representing Southern hemisphere destinations) at four travel-relevant dates.

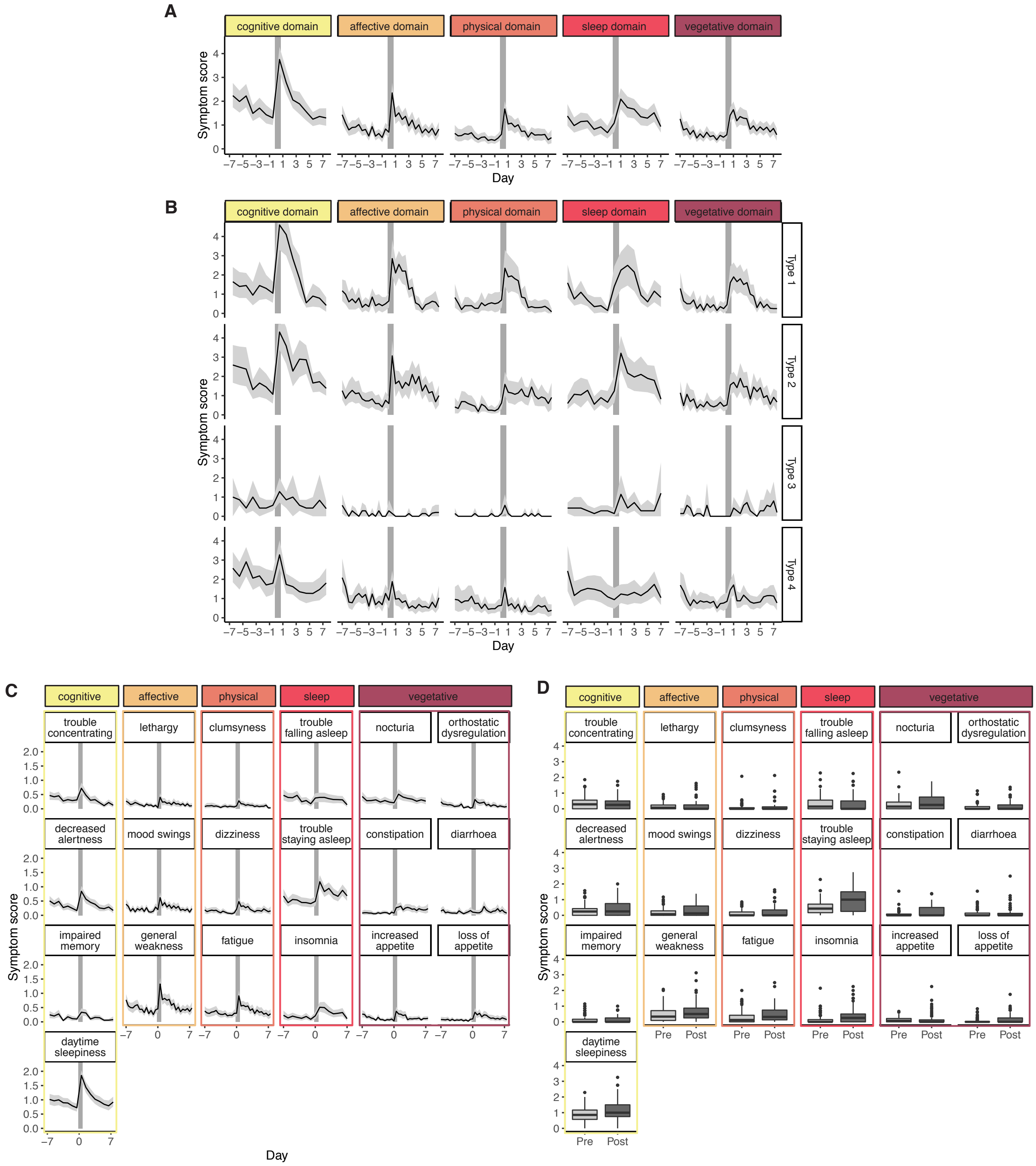


**Figure S2. Scores for individual symptom domains and symptoms. Related to Figure 1.** Scores from the Charité Jetlag Scale in all 90 participants reported twice daily for 7 days before and after the flight. Trajectories of mean symptom score (± 95% CI) per time point A) for each of the 5 symptom domains, B) per trajectory type and symptom domain, C) for each of the 19 individual symptoms. Grey bars indicate the day of flight. D) Comparison of individual mean symptom scores for all 19 individual symptoms between all days before the flight and 4 days thereafter.

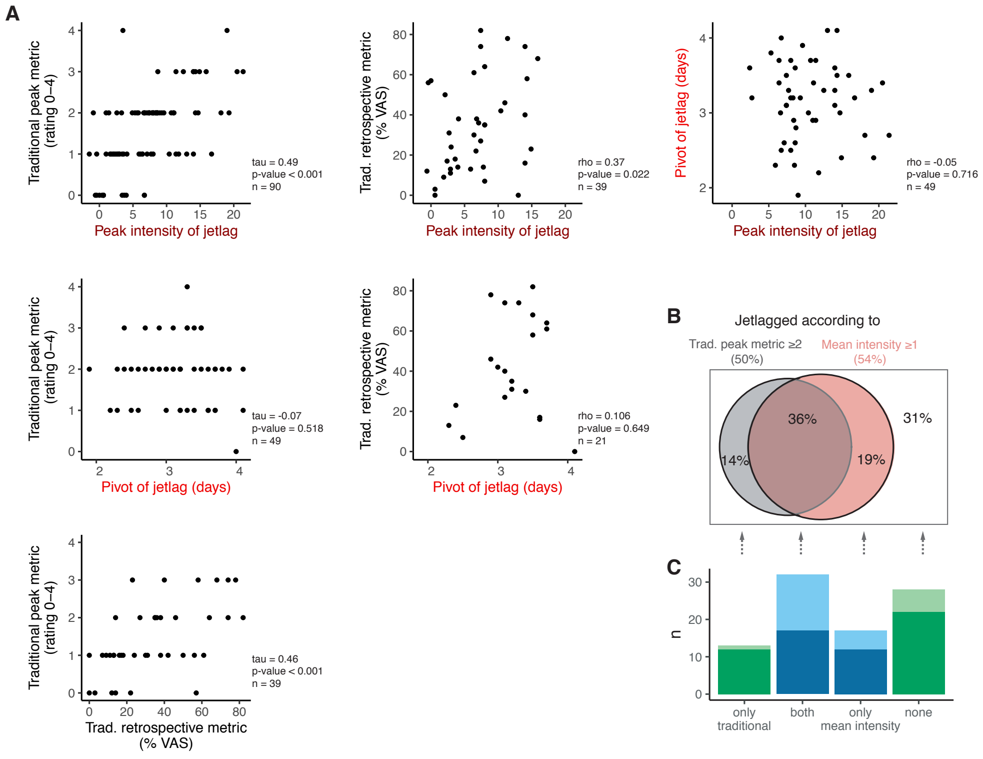


**Figure S3. Relationship among jetlag measures. Related to Figure 2.**
A) Further relationships among indicated jetlag measures complementing those depicted in Figure 2C. Results of correlation analyses (Spearman rho, Kendall tau) are provided. B) Venn diagram indicating proportion of sample classified as jetlagged according to two intra-individual split criterions on mean intensity and traditional peak metric. C) The number of participants for each symptom trajectory type underlying the different areas from B.

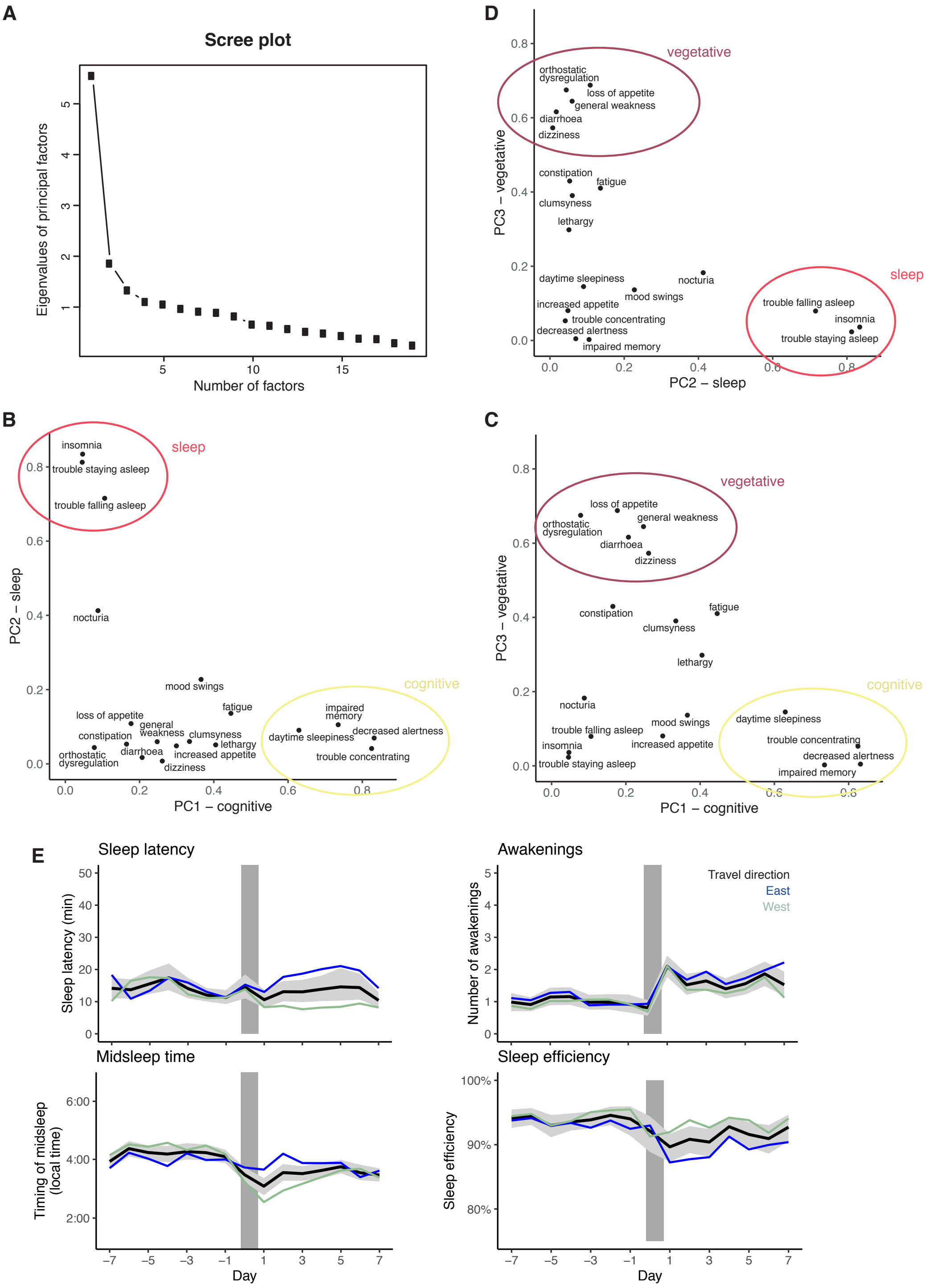


**Figure S4. Overview of alternative jetlag outcomes. Related to Figure 3 and Tables S3 and S4.** A-D) Principal component analysis of daily symptom scores for all 19 individual symptoms from the Charité Jetlag Scale (n=1295 participant days) performed for optimal data reduction. A) Scree plot suggesting the use of 3 principal components (PC1-3). B-D) Scatterplots showing the loading of each individual symptom on each principal component. The symptoms with the highest loadings for each component are circled. E) Trajectories of sleep outcomes from daily sleep diaries depicting effects detected in regression analyses (Tab. S5). Plotted are the overall mean (black) and 95% CI (grey) together with means for both travel directions (blue, East; green, West). Grey bars indicate day of flight. Note that the y-axis for midsleep time is scaled in local time, i.e. for the pre-flight period the time at origin, for the post-flight period the time at destination. Therefore, the small change in midsleep relative to the number of time zones crossed indicates an almost instantaneous resetting of sleep timing post-flight.

**Table S1. Results of logistic regression analysis for post-flight status. Related to Figure 1.**
To identify the symptom domains whose increase is most indicative of a post-flight status and thus potentially most indicative of jetlag disorder, logistic regression analysis was performed on individual domain scores. For maximum information, individual scores for each domain and questionnaire instance were entered rather than pre-/post-flight aggregates, thus mixed effects-model were calculated to account for the repeated measures nature of the data. Note, however, the absence of any variance in ID ($T_{00}$ record_id). Bold print marks p-values below the alpha level of 0.05. Abbreviations: CI, 95% confidence intervals; p, p-value; $\sigma^2$, variance of residuals of random effects; $\tau_{00}$, variance of ID intercepts of random effects; N, number of participants; Marginal $R^2$ describes the amount of variance explained by the fixed effects (predictors).

| | Post-flight status | | |
|---|---|---|---|
| *Predictors* | *Odds Ratio* | *CI* | *p* |
| (Intercept) | 0.74 | 0.62 – 0.88 | **0.001** |
| Sleep domain | 1.16 | 1.07 – 1.24 | **<0.001** |
| Cognitive domain | 0.88 | 0.82 – 0.96 | **0.002** |
| Affective domain | 1.00 | 0.88 – 1.17 | 0.932 |
| Physical domain | 1.05 | 0.96 – 1.33 | 0.223 |
| Vegetative domain | 1.17 | 1.14 – 1.44 | **<0.001** |
| **Random Effects** | | | |
| $\sigma^2$ | 3.29 | | |
| $T_{00\ record\_id}$ | 0.00 | | |
| $N_{record\_id}$ | 90 | | |
| Observations | 1206 | | |
| Marginal $R^2$ / Conditional $R^2$ | 0.066 / NA | | |

**Table S2. Results of regression analyses for main jetlag outcomes with expected jetlag duration as predictor. Related to Figure 3.**
Linear regression results for the 4 main jetlag outcomes peak intensity, mean intensity, pivot and traditional peak metric. Results for both standardized outcomes (better comparability, as in Fig. 3B) and unstandardized outcomes (easier interpretation) are provided. In addition, for peak intensity, both the results for the log-transformed outcome (fulfilling heteroscedacity assumption, $\log_{10}(x+3)$) and the untransformed outcome (easier interpretation, as in Fig. 3B) are listed. Bold print marks p-values below the original alpha level of 0.05, but multiple testing (8 models, 4 outcomes x 2 expectation measures) leads to an adjusted alpha level of 0.00625 (grey shading). Abbreviations: Est., estimates; CI, 95% confidence intervals; p, p-value; DF, degrees of freedom; $R^2$ adjusted is the explanatory power accounting for the number of predictors in the model.

| | Peak Intensity | | | Peak Intensity (log-transformed) | | | Mean Intensity | | | Pivot of Jetlag | | | Traditional Peak Metric | | |
|---|---|---|---|---|---|---|---|---|---|---|---|---|---|---|---|
| *Predictors* | *Est.* | *CI* | *p* | *Est.* | *CI* | *p* | *Est.* | *CI* | *p* | *Est.* | *CI* | *p* | *Est.* | *CI* | *p* |
| **Standardized Outcomes** | | | | | | | | | | | | | | | |
| (Intercept) | -0.36 | -2.52 – 1.80 | 0.742 | -0.49 | -2.66 – 1.67 | 0.651 | 0.01 | -2.37 – 2.39 | 0.995 | -0.15 | -3.42 – 3.12 | 0.927 | 0.48 | -1.66 – 2.63 | 0.655 |
| Age (y) | -0.00 | -0.06 – 0.05 | 0.958 | -0.01 | -0.06 – 0.05 | 0.839 | -0.01 | -0.07 – 0.05 | 0.776 | -0.01 | -0.09 – 0.08 | 0.893 | 0.03 | -0.03 – 0.08 | 0.358 |
| Gender [female] | 0.30 | -0.14 – 0.75 | 0.181 | 0.27 | -0.18 – 0.72 | 0.236 | -0.03 | -0.53 – 0.47 | 0.916 | 0.10 | -0.58 – 0.79 | 0.764 | 0.08 | -0.37 – 0.52 | 0.736 |
| Health (1/10 VAS) | -0.02 | -0.17 – 0.13 | 0.767 | 0.01 | -0.14 – 0.16 | 0.867 | 0.01 | -0.16 – 0.17 | 0.926 | 0.11 | -0.13 – 0.35 | 0.370 | -0.22 | -0.37 – -0.07 | **0.004** |
| Chronotype (h, centred) | 0.22 | -0.06 – 0.50 | 0.125 | 0.19 | -0.09 – 0.48 | 0.176 | 0.08 | -0.24 – 0.39 | 0.625 | -0.02 | -0.42 – 0.39 | 0.938 | 0.16 | -0.13 – 0.44 | 0.273 |
| Direction [west] | -0.27 | -0.67 – 0.13 | 0.181 | -0.30 | -0.70 – 0.09 | 0.132 | -0.25 | -0.69 – 0.19 | 0.265 | 0.07 | -0.57 – 0.71 | 0.822 | 0.04 | -0.35 – 0.44 | 0.832 |
| Chronotype*Direction [west] | -0.31 | -0.72 – 0.11 | 0.144 | -0.43 | -0.84 – -0.02 | **0.042** | -0.32 | -0.79 – 0.14 | 0.170 | 0.13 | -0.62 – 0.88 | 0.720 | -0.33 | -0.74 – 0.08 | 0.112 |
| Time zones crossed | -0.02 | -0.14 – 0.09 | 0.664 | -0.02 | -0.13 – 0.10 | 0.762 | -0.01 | -0.14 – 0.11 | 0.845 | -0.15 | -0.33 – 0.03 | 0.093 | 0.02 | -0.09 – 0.14 | 0.687 |
| Expected jetlag duration (days) | 0.24 | 0.13 – 0.34 | **<0.001** | 0.22 | 0.12 – 0.33 | **<0.001** | 0.13 | 0.01 – 0.24 | **0.028** | 0.08 | -0.06 – 0.22 | 0.277 | 0.17 | 0.06 – 0.27 | **0.002** |
| **Unstandardized Outcomes** | | | | | | | | | | | | | | | |
| (Intercept) | 5.25 | -6.33 – 16.83 | 0.370 | 0.81 | 0.24 – 1.37 | 0.006 | 1.56 | -4.63 – 7.74 | 0.618 | 3.07 | 1.35 – 4.79 | 0.001 | 2.01 | -0.00 – 4.02 | 0.050 |
| Age (y) | -0.01 | -0.31 – 0.29 | 0.958 | -0.00 | -0.02 – 0.01 | 0.839 | -0.02 | -0.18 – 0.14 | 0.776 | -0.00 | -0.05 – 0.04 | 0.893 | 0.02 | -0.03 – 0.08 | 0.358 |
| Gender [female] | 1.63 | -0.77 – 4.03 | 0.181 | 0.07 | -0.05 – 0.19 | 0.236 | -0.07 | -1.37 – 1.23 | 0.916 | 0.05 | -0.31 – 0.42 | 0.764 | 0.07 | -0.35 – 0.49 | 0.736 |
| Health (1/10 VAS) | -0.12 | -0.92 – 0.68 | 0.767 | 0.00 | -0.04 – 0.04 | 0.867 | 0.02 | -0.41 – 0.45 | 0.926 | 0.06 | -0.07 – 0.18 | 0.370 | -0.21 | -0.34 – -0.07 | **0.004** |
| Chronotype (h, centred) | 1.18 | -0.33 – 2.70 | 0.125 | 0.05 | -0.02 – 0.12 | 0.176 | 0.20 | -0.61 – 1.01 | 0.625 | -0.01 | -0.22 – 0.20 | 0.938 | 0.15 | -0.12 – 0.41 | 0.273 |
| Direction [west] | -1.44 | -3.57 – 0.69 | 0.181 | -0.08 | -0.18 – 0.02 | 0.132 | -0.65 | -1.79 – 0.50 | 0.265 | 0.04 | -0.30 – 0.37 | 0.822 | 0.04 | -0.33 – 0.41 | 0.832 |
| Chronotype*Direction [west] | -1.64 | -3.86 – 0.57 | 0.144 | -0.11 | -0.22 – -0.00 | **0.042** | -0.84 | -2.05 – 0.37 | 0.170 | 0.07 | -0.32 – 0.46 | 0.720 | -0.31 | -0.70 – 0.07 | 0.112 |
| Time zones crossed | -0.13 | -0.74 – 0.48 | 0.664 | -0.00 | -0.03 – 0.03 | 0.762 | -0.03 | -0.36 – 0.30 | 0.845 | -0.08 | -0.17 – 0.01 | 0.093 | 0.02 | -0.08 – 0.13 | 0.687 |
| Expected jetlag duration (days) | 1.27 | 0.71 – 1.83 | **<0.001** | 0.06 | 0.03 – 0.08 | **<0.001** | 0.34 | 0.04 – 0.64 | **0.028** | 0.04 | -0.03 – 0.11 | 0.277 | 0.16 | 0.06 – 0.25 | **0.002** |
| Observations | 90 | | | 90 | | | 89 | | | 49 | | | 90 | | |
| $R^2$/$R^2$ adjusted | 0.130 / 0.045 | | | 0.160 / 0.077 | | | 0.073 / -0.019 | | | 0.084 / -0.099 | | | 0.239 / 0.163 | | |
| F-statistic | 1.519 | | | 1.93 | | | 0.793 | | | 0.4593 | | | 3.174 | | |
| DF | 8 / 81 | | | 8 / 81 | | | 8 / 80 | | | 8 / 40 | | | 8 / 81 | | |
| p-value | 0.1636 | | | 0.0664 | | | 0.6103 | | | 0.8771 | | | 0.003525 | | |

Reference category for Gender is male and for Direction east.

**Table S3. Results of regression analyses for main jetlag outcomes with expected jetlag intensity as predictor. Related to Figure 3.**
Linear regression results for the 4 main jetlag outcomes peak intensity, mean intensity, pivot and traditional peak metric. Results for both standardized outcomes (better comparability, as in Fig. 3B) and unstandardized outcomes (easier interpretation) are provided. In addition, for peak intensity, both the results for the log-transformed outcome (fulfilling heteroscedacity assumption, $\log_{10}(x+3)$) and the untransformed outcome (easier interpretation, as in Fig. 3B) are listed. Bold print mark p-values below the original alpha level of 0.05, but multiple testing (8 models, 4 outcomes x 2 expectation measures) leads to an adjusted alpha level of 0.00625 (grey shading). Abbreviations: Est., estimates; CI, 95% confidence intervals; p, p-value; DF, degrees of freedom; $R^2$ adjusted is the explanatory power accounting for the number of predictors in the model.

| | Peak Intensity | | | Peak Intensity (log-transformed) | | | Mean Intensity | | | Pivot of Jetlag | | | Traditional Peak Metric | | |
|---|---|---|---|---|---|---|---|---|---|---|---|---|---|---|---|
| *Predictors* | *Est.* | *CI* | *p* | *Est.* | *CI* | *p* | *Est.* | *CI* | *p* | *Est.* | *CI* | *p* | *Est.* | *CI* | *p* |
| **Standardized Outcomes** | | | | | | | | | | | | | | | |
| (Intercept) | -0.18 | -2.54 – 2.18 | 0.878 | -0.38 | -2.69 – 1.94 | 0.748 | 0.10 | -2.34 – 2.53 | 0.938 | -0.05 | -3.36 – 3.26 | 0.975 | 0.51 | -1.70 – 2.72 | 0.646 |
| Age (y) | 0.01 | -0.05 – 0.07 | 0.810 | 0.00 | -0.06 – 0.06 | 0.986 | -0.00 | -0.07 – 0.06 | 0.886 | -0.00 | -0.09 – 0.09 | 0.979 | 0.03 | -0.03 – 0.09 | 0.338 |
| Gender [female] | 0.11 | -0.38 – 0.60 | 0.647 | 0.08 | -0.40 – 0.56 | 0.740 | -0.14 | -0.65 – 0.37 | 0.591 | 0.01 | -0.72 – 0.74 | 0.978 | -0.08 | -0.54 – 0.38 | 0.724 |
| Health (1/10 VAS) | -0.04 | -0.20 – 0.12 | 0.623 | -0.00 | -0.16 – 0.16 | 0.999 | -0.00 | -0.17 – 0.17 | 0.990 | 0.10 | -0.15 – 0.35 | 0.421 | -0.22 | -0.38 – -0.07 | **0.005** |
| Chronotype (h, centred) | 0.19 | -0.12 – 0.50 | 0.221 | 0.18 | -0.13 – 0.49 | 0.246 | 0.06 | -0.26 – 0.39 | 0.693 | -0.02 | -0.46 – 0.42 | 0.916 | 0.16 | -0.13 – 0.45 | 0.279 |
| Direction [west] | -0.32 | -0.75 – 0.12 | 0.151 | -0.34 | -0.76 – 0.09 | 0.121 | -0.27 | -0.73 – 0.18 | 0.230 | 0.05 | -0.59 – 0.70 | 0.866 | 0.04 | -0.37 – 0.44 | 0.863 |
| Chronotype*Direction [west] | -0.30 | -0.75 – 0.15 | 0.192 | -0.43 | -0.87 – 0.02 | 0.059 | -0.33 | -0.80 – 0.15 | 0.176 | 0.15 | -0.62 – 0.92 | 0.697 | -0.33 | -0.76 – 0.09 | 0.119 |
| Time zones crossed | -0.01 | -0.13 – 0.12 | 0.924 | -0.00 | -0.13 – 0.12 | 0.956 | -0.00 | -0.13 – 0.13 | 0.973 | -0.15 | -0.34 – 0.04 | 0.130 | 0.03 | -0.09 – 0.14 | 0.627 |
| Expected jetlag intensity (1/10 VAS) | 0.10 | 0.00 – 0.21 | **0.046** | 0.11 | 0.01 – 0.21 | **0.025** | 0.06 | -0.04 – 0.17 | 0.254 | 0.03 | -0.15 – 0.21 | 0.730 | 0.11 | 0.02 – 0.21 | **0.023** |
| **Unstandardized Outcomes** | | | | | | | | | | | | | | | |
| (Intercept) | 6.19 | -6.47 – 18.84 | 0.333 | 0.84 | 0.24 – 1.45 | **0.007** | 1.79 | -4.55 – 8.12 | 0.576 | 3.12 | 1.38 – 4.86 | **0.001** | 2.03 | -0.03 – 4.10 | 0.054 |
| Age (y) | 0.04 | -0.29 – 0.37 | 0.810 | 0.00 | -0.02 – 0.02 | 0.986 | -0.01 | -0.18 – 0.15 | 0.886 | -0.00 | -0.05 – 0.05 | 0.979 | 0.03 | -0.03 – 0.08 | 0.338 |
| Gender [female] | 0.61 | -2.01 – 3.23 | 0.647 | 0.02 | -0.10 – 0.15 | 0.740 | -0.36 | -1.69 – 0.97 | 0.591 | 0.01 | -0.38 – 0.39 | 0.978 | -0.08 | -0.50 – 0.35 | 0.724 |
| Health (1/10 VAS) | -0.22 | -1.10 – 0.66 | 0.623 | -0.00 | -0.04 – 0.04 | 0.999 | -0.00 | -0.44 – 0.44 | 0.990 | 0.05 | -0.08 – 0.18 | 0.421 | -0.21 | -0.35 – -0.07 | **0.005** |
| Chronotype (h, centred) | 1.04 | -0.64 – 2.71 | 0.221 | 0.05 | -0.03 – 0.13 | 0.246 | 0.17 | -0.67 – 1.00 | 0.693 | -0.01 | -0.24 – 0.22 | 0.916 | 0.15 | -0.12 – 0.42 | 0.279 |
| Direction [west] | -1.70 | -4.03 – 0.63 | 0.151 | -0.09 | -0.20 – 0.02 | 0.121 | -0.71 | -1.89 – 0.46 | 0.230 | 0.03 | -0.31 – 0.37 | 0.866 | 0.03 | -0.35 – 0.41 | 0.863 |
| Chronotype*Direction [west] | -1.60 | -4.02 – 0.82 | 0.192 | -0.11 | -0.23 – 0.00 | 0.059 | -0.85 | -2.08 – 0.39 | 0.176 | 0.08 | -0.33 – 0.48 | 0.697 | -0.31 | -0.71 – 0.08 | 0.119 |
| Time zones crossed | -0.03 | -0.70 – 0.64 | 0.924 | -0.00 | -0.03 – 0.03 | 0.956 | -0.01 | -0.34 – 0.33 | 0.973 | -0.08 | -0.18 – 0.02 | 0.130 | 0.03 | -0.08 – 0.14 | 0.627 |
| Expected jetlag intensity (1/10 VAS) | 0.56 | 0.01 – 1.10 | **0.046** | 0.03 | 0.00 – 0.06 | **0.025** | 0.16 | -0.12 – 0.43 | 0.254 | 0.02 | -0.08 – 0.11 | 0.730 | 0.10 | 0.01 – 0.19 | **0.023** |
| Observations | | 90 | | | 90 | | | 89 | | | 49 | | | 90 | |
| $R^2$/$R^2$ adjusted | | 0.130 / 0.045 | | | 0.160 / 0.077 | | | 0.073 / -0.019 | | | 0.084 / -0.099 | | | 0.239 / 0.163 | |
| F-statistic | | 1.519 | | | 1.93 | | | 0.793 | | | 0.4593 | | | 3.174 | |
| DF | | 8 / 81 | | | 8 / 81 | | | 8 / 80 | | | 8 / 40 | | | 8 / 81 | |
| p-value | | 0.1636 | | | 0.0664 | | | 0.6103 | | | 0.8771 | | | 0.003525 | |

Reference category for Gender is male and for Direction east.

**Table S4. Results of regression analyses for alternative jetlag outcomes.**
Bold print marks p-values below the original alpha level of 0.05, but multiple testing (9 models) leads to an adjusted alpha level of 0.0056 (grey shading). Abbreviations: Est., Estimates; CI, 95% confidence intervals; p, p-value; DF, degrees of freedom; OR, odds ratios; PC, principal component

| | **PC1 Cognitive symptoms** | | | **PC2 Sleep symptoms** | | | **PC3 Vegetative symptoms** | | | **Classified as jetlagged** | | |
|---|---|---|---|---|---|---|---|---|---|---|---|---|
| *Predictors* | *Est.* | *CI* | *p* | *Est.* | *CI* | *p* | *Est.* | *CI* | *p* | *OR* | *CI* | *p* |
| (Intercept) | 0.24 | -1.53 – 2.01 | 0.786 | 0.19 | -1.99 – 2.36 | 0.864 | 0.71 | -0.77 – 2.18 | 0.342 | 0.01 | 0.00 – 1.31 | 0.070 |
| Age (y) | -0.00 | -0.05 – 0.04 | 0.886 | -0.00 | -0.06 – 0.05 | 0.866 | -0.01 | -0.05 – 0.03 | 0.574 | 1.05 | 0.92 – 1.21 | 0.439 |
| Gender [female] | 0.04 | -0.33 – 0.41 | 0.829 | 0.19 | -0.26 – 0.65 | 0.394 | -0.08 | -0.39 – 0.23 | 0.611 | 1.16 | 0.39 – 3.43 | 0.791 |
| Health (1/10 VAS) | -0.01 | -0.13 – 0.12 | 0.923 | -0.01 | -0.16 – 0.14 | 0.858 | -0.01 | -0.11 – 0.09 | 0.874 | 1.46 | 1.00 – 2.20 | 0.060 |
| Chronotype (h, centred) | -0.17 | -0.40 – 0.06 | 0.145 | 0.18 | -0.10 – 0.47 | 0.207 | 0.21 | 0.02 – 0.40 | **0.034** | 1.42 | 0.73 – 2.91 | 0.313 |
| Direction [west] | 0.12 | -0.21 – 0.44 | 0.474 | -0.34 | -0.74 – 0.06 | 0.093 | -0.21 | -0.48 – 0.06 | 0.127 | 0.78 | 0.30 – 2.02 | 0.609 |
| Chronotype*Direction [west] | 0.31 | -0.03 – 0.65 | 0.071 | -0.29 | -0.70 – 0.13 | 0.176 | -0.45 | -0.74 – -0.16 | **0.002** | 0.60 | 0.21 – 1.59 | 0.316 |
| Time zones crossed | -0.06 | -0.15 – 0.04 | 0.233 | 0.04 | -0.07 – 0.16 | 0.468 | 0.00 | -0.08 – 0.08 | 0.975 | 0.97 | 0.73 – 1.28 | 0.822 |
| Expected jetlag duration (d) | 0.08 | -0.01 – 0.16 | 0.084 | 0.09 | -0.01 – 0.20 | 0.088 | 0.06 | -0.01 – 0.13 | 0.096 | 1.61 | 1.12 – 2.49 | **0.020** |
| Observations | 90 | | | 90 | | | 89 | | | 90 | | |
| $R^2/R^2$ adjusted | 0.090 / 0.001 | | | 0.120 / 0.033 | | | 0.167 / 0.084 | | | 0.119 | | |
| F-statistic | 1.006 | | | 1.382 | | | 2.012 | | | | | |
| DF | 8 / 81 | | | 8 / 81 | | | 8/80 | | | | | |
| p-value | 0.4384 | | | 0.2168 | | | 0.0553 | | | | | |

| | **Cognitive domain** | | | **Affective domain (log-transformed)** | | | **Physical domain** | | | **Vegetative domain** | | | **Sleep domain** | | |
|---|---|---|---|---|---|---|---|---|---|---|---|---|---|---|---|
| *Predictors* | *Est.* | *CI* | *p* | *Est.* | *CI* | *p* | *Est.* | *CI* | *p* | *Est.* | *CI* | *p* | *Est.* | *CI* | *p* |
| (Intercept) | 0.34 | -3.28 – 3.95 | 0.854 | 0.53 | 0.24 – 0.82 | **0.001** | 1.20 | -0.41 – 2.81 | 0.142 | -1.36 | -3.26 – 0.54 | 0.159 | -0.09 | -3.90 – 3.71 | 0.961 |
| Age (y) | -0.00 | -0.09 – 0.09 | 0.993 | -0.00 | -0.01 – 0.01 | 0.681 | -0.03 | -0.07 – 0.01 | 0.205 | 0.03 | -0.02 – 0.08 | 0.214 | -0.00 | -0.10 – 0.10 | 0.981 |
| Gender [female] | 0.10 | -0.65 – 0.85 | 0.786 | 0.00 | -0.06 – 0.06 | 0.943 | -0.06 | -0.40 – 0.27 | 0.705 | -0.01 | -0.40 – 0.38 | 0.978 | 0.41 | -0.38 – 1.19 | 0.310 |
| Health (1/10 VAS) | -0.01 | -0.26 – 0.24 | 0.957 | 0.00 | -0.02 – 0.02 | 0.669 | -0.05 | -0.17 – 0.06 | 0.351 | 0.06 | -0.07 – 0.20 | 0.338 | 0.02 | -0.25 – 0.28 | 0.908 |
| Chronotype (h, centred) | -0.31 | -0.78 – 0.16 | 0.196 | -0.02 | -0.05 – 0.02 | 0.428 | 0.24 | 0.03 – 0.45 | **0.028** | 0.06 | -0.18 – 0.30 | 0.628 | 0.41 | -0.09 – 0.91 | 0.104 |
| Direction [west] | 0.20 | -0.47 – 0.86 | 0.555 | 0.00 | -0.05 – 0.06 | 0.960 | -0.23 | -0.53 – 0.07 | 0.128 | 0.02 | -0.32 – 0.36 | 0.903 | -0.77 | -1.47 – -0.07 | **0.031** |
| Chronotype*Direction [west] | 0.49 | -0.20 – 1.18 | 0.163 | 0.02 | -0.03 – 0.08 | 0.421 | -0.45 | -0.76 – -0.13 | **0.006** | -0.40 | -0.76 – -0.03 | **0.032** | -0.60 | -1.33 – 0.13 | 0.104 |
| Time zones crossed | -0.09 | -0.28 – 0.10 | 0.333 | -0.01 | -0.02 – 0.01 | 0.277 | -0.01 | -0.09 – 0.08 | 0.874 | 0.08 | -0.01 – 0.18 | 0.089 | 0.07 | -0.13 – 0.27 | 0.508 |
| Expected jetlag duration (d) | 0.14 | -0.03 – 0.31 | 0.115 | 0.01 | -0.00 – 0.02 | 0.148 | 0.13 | 0.06 – 0.21 | **0.001** | 0.02 | -0.07 – 0.11 | 0.666 | 0.15 | -0.04 – 0.33 | 0.119 |
| Observations | 90 | | | 90 | | | 87 | | | 88 | | | 90 | | |
| $R^2/R^2$ adjusted | 0.069 / -0.023 | | | 0.051 / -0.042 | | | 0.247 / 0.170 | | | 0.140 / 0.053 | | | 0.142 / 0.057 | | |
| F-statistic | 0.7462 | | | 0.5465 | | | 3.204 | | | 1.605 | | | 1.673 | | |
| DF | 8 / 81 | | | 8 / 81 | | | 8 / 78 | | | 8 / 79 | | | 8 / 81 | | |
| p-value | 0.6506 | | | 0.818 | | | 0.0034 | | | 0.1366 | | | 0.1176 | | |

Reference category for Gender is male and for Direction east.

**Table S5. Results of regression analyses for sleep outcomes.**

Linear regression results for sleep outcomes originating from daily sleep diary. Specifically, outcomes were changes (Δ) in each listed sleep parameter from individual baseline mean to individual 4-day-post-flight mean analogous to the calculation of the mean intensity of jetlag. Bold print marks p-values below the original alpha level of 0.05, but multiple testing (4 models) leads to an adjusted alpha level of 0.0125 (grey shading). Abbreviations: Est., estimates; CI, 95% confidence intervals; p, p-value; DF, degrees of freedom; $R^2$ adjusted is the explanatory power accounting for the number of predictors in the model.

| | **Δ Awakenings (n)** | | | **Δ Midsleep (h)** | | | **Δ Sleep latency (min)** | | | **Δ Sleep efficiency (%)** | | |
|---|---|---|---|---|---|---|---|---|---|---|---|---|
| *Predictors* | *Est.* | *CI* | *p* | *Est.* | *CI* | *p* | *Est.* | *CI* | *p* | *Est.* | *CI* | *p* |
| (Intercept) | 0.70 | -1.70 – 3.10 | 0.562 | -1.90 | -4.22 – 0.42 | 0.107 | -17.24 | -44.55 – 10.08 | 0.213 | 10.54 | -2.45 – 23.53 | 0.110 |
| Age (y) | -0.03 | -0.09 – 0.03 | 0.366 | 0.01 | -0.05 – 0.07 | 0.639 | 0.27 | -0.44 – 0.98 | 0.455 | -0.27 | -0.60 – 0.07 | 0.114 |
| Gender [female] | -0.02 | -0.52 – 0.48 | 0.941 | 0.31 | -0.17 – 0.79 | 0.201 | -0.01 | -5.53 – 5.51 | 0.998 | -2.45 | -5.13 – 0.23 | 0.073 |
| Health (1/10 VAS) | -0.01 | -0.18 – 0.16 | 0.901 | 0.24 | 0.08 – 0.40 | **0.004** | 1.62 | -0.21 – 3.45 | 0.083 | -0.65 | -1.56 – 0.26 | 0.160 |
| Chronotype | -0.05 | -0.37 – 0.26 | 0.740 | -0.06 | -0.37 – 0.24 | 0.685 | -2.07 | -5.77 – 1.64 | 0.270 | -2.16 | -3.93 – -0.39 | **0.018** |
| Direction [west] | -0.22 | -0.66 – 0.22 | 0.331 | -1.32 | -1.75 – -0.90 | **<0.001** | -9.24 | -13.94 – -4.55 | **<0.001** | 3.21 | 0.84 – 5.59 | **0.009** |
| Chronotype*Direction [west] | -0.15 | -0.61 – 0.31 | 0.532 | -0.61 | -1.05 – -0.17 | **0.008** | 3.54 | -1.61 – 8.69 | 0.175 | 1.14 | -1.39 – 3.67 | 0.372 |
| Time zones crossed | 0.12 | -0.00 – 0.25 | 0.055 | -0.06 | -0.18 – 0.06 | 0.337 | 0.07 | -1.27 – 1.41 | 0.915 | -0.12 | -0.81 – 0.56 | 0.719 |
| Expected jetlag duration (d) | 0.02 | -0.10 – 0.14 | 0.732 | -0.10 | -0.21 – 0.01 | 0.086 | 0.23 | -1.09 – 1.54 | 0.733 | -0.23 | -0.86 – 0.39 | 0.459 |
| Observations | 90 | | | 90 | | | 85 | | | 88 | | |
| $R^2$/$R^2$ adjusted | 0.077 / -0.014 | | | 0.502 / 0.453 | | | 0.216 / 0.133 | | | 0.170 / 0.086 | | |
| F-statistic | 0.8437 | | | 10.21 | | | 2.61 | | | 2.021 | | |
| DF | 8 / 81 | | | 8 / 81 | | | 8 / 76 | | | 8 / 79 | | |
| p-value | 0.5672 | | | <0.0001 | | | 0.0141 | | | 0.0544 | | |

Reference category for Gender is male and for Direction east.
